# A summary of intraspecific size variation for large mysticetes

**DOI:** 10.1101/2025.02.07.637173

**Authors:** Joseph McClure

**Affiliations:** N/A

**Keywords:** Baleen whales, Mysticetes, Cetaceans, Body size, Body mass, Intraspecific variation

## Abstract

Following existing work on odontocetes, an extensive survey of literature was conducted to summarize the growth and body mass of nine mysticete species along with analyses of adult size distributions using data from the International Whaling Commission catch database. Relationships uncovered between length parameters in mysticetes are combined with results previously obtained for odontocetes. On average, the mean adult length was 4% below the asymptotic size and its relative variability appeared to be very similar across most species (CV=4-5%). This value was expectedly higher in species with more prolonged growth throughout adulthood (CV=6-8%). Total body mass relationships between mysticete species appear to align with feeding strategies, as generalists and skim-feeders are notably heavier than lunge-feeding specialists.

## 1 Introduction

Knowledge on adult size variation in cetaceans can have various applications researching their ecology (Pershing et al., 2010), conservation (Christiansen et al., 2020), stock divisions (Best, 1977; Ichihara, 1966; Pastene et al., 2020), and evolution (Bianucci et al., 2019; Bisconti et al., 2021). Due to the broader focus across multiple taxa, one review was only able to examine the literature and commercial data for *Balaenoptera musculus* (McClain et al., 2015). Since then, independent advances were made for other species (Agbayani et al., 2020; Christiansen et al., 2022; Fortune et al., 2021; George et al., 2021; Russell et al., 2023), warranting a deeper review of existing patterns in adult size variation and body mass relationships.

A comprehensive effort was placed into reviewing the adult variation in total body length (TBL) and body mass relationships of large odontocetes (McClure, 2024). However, the limited number of sampled species prevented strong conclusions. The previous analysis will now be extended towards nine mysticete species spanning five genera. The mysticetes selected include all extant members of the Balaenidae family, *Eschrichtius robustus* (Lilljeborg, 1861), *Megaptera novaeangliae* (Borowski, 1781), and the *Balaenoptera* species ranging in size from *B. edeni* (Anderson, 1878) to *B. musculus* (Linneaus, 1758).

In general, large baleen whales are weaned within one year, sexually mature after 5-10 years, and physically mature after 25-40 years (Agbayani et al., 2020; Aguilar & Lockyer, 1987; Bando, 2021; Best, 2011; Best & Lockyer, 2002; Branch, 2008; Chittleborough, 1965; Fortune et al., 2021). In *Balaena mysticetus* (Linneaus, 1758), sexual maturity occurs after 25 years and physical growth continues much further into senescence (George et al., 1999; Tarpley et al., 2021). As done in the previous analysis (McClure, 2024), the correlation between the length interval from sexual to physical maturity and the coefficients of variation (CV) in mean adult size will be examined for each species.

An unprecedented review of existing literature was conducted along with an analysis of length distributions within the IWC catch database. Analyses on body mass relationships, largest reliable sizes, and postnatal growth are also provided. The data presented in this review will assist future ecological, anatomical, energetic, and phylogenetic research that benefits from incorporating intraspecific information about TBL and body mass.

## 2 Methods

The nine mysticete species under review are summarized in Table 1. Literature was comprehensively surveyed for morphometric data and growth parameters using the exact standards described in the previous review for odontocetes (McClure, 2024). Data on the mean TBL at birth (*L*_b_), weaning (*L*_w_), sexual maturity (*L*_sm_), and physical maturity (*L*^∞^) are summarized here. In total, 340 sources from literature (73 books, 26 conference papers, 229 published articles, 8 technical reports/research documents, 3 theses, and 1 media article) were surveyed and 126 were selected after filtering.

**Table 1:**
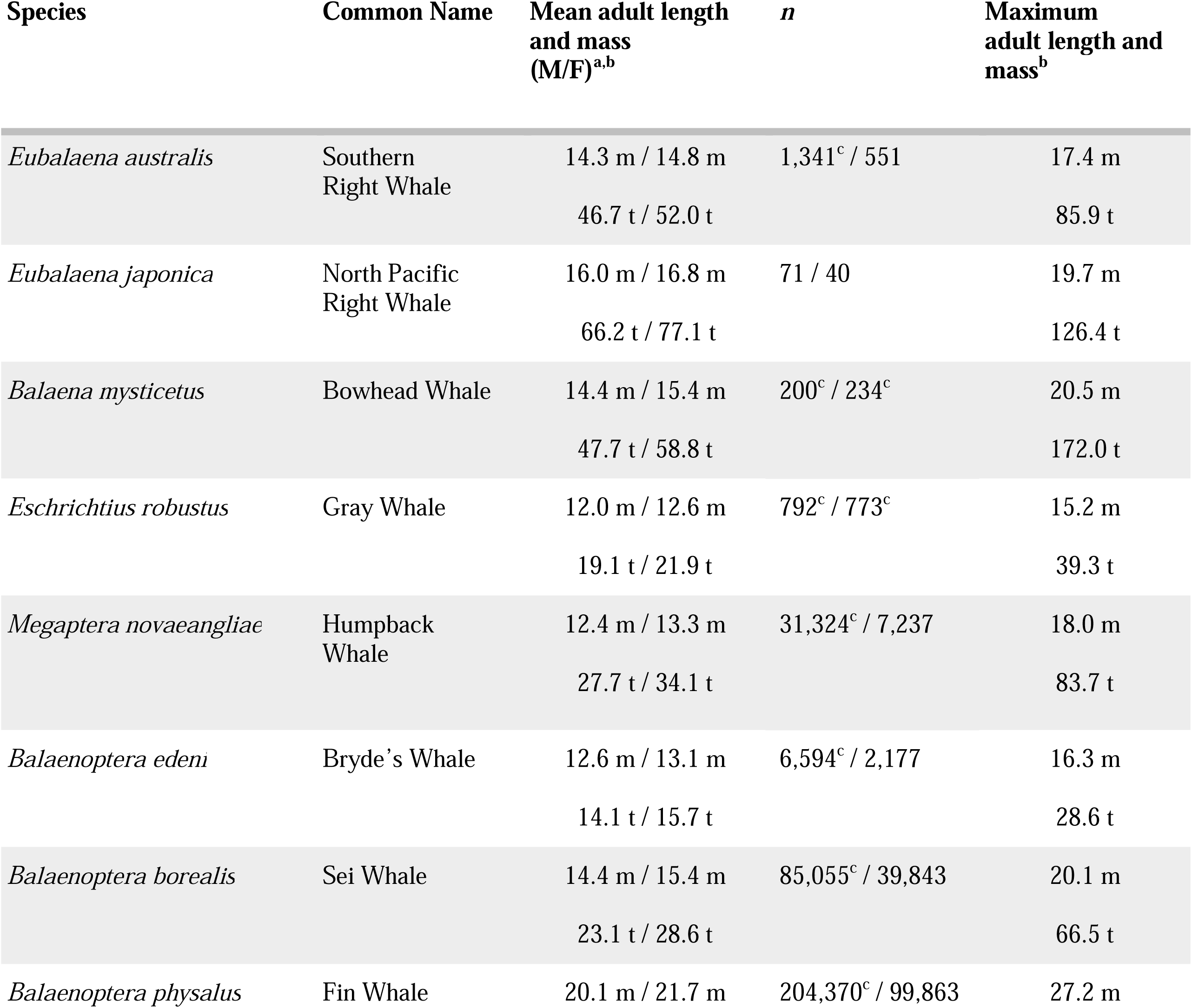
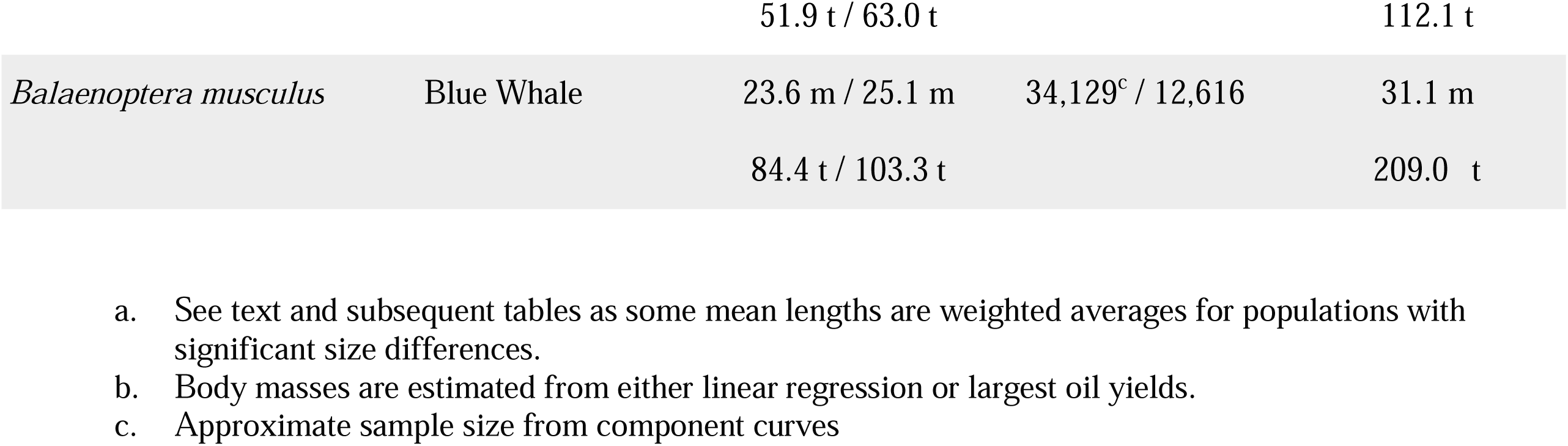
Summary of adult body length and mass for nine mysticete species.

| Species | Common Name | Mean adult length and mass (M/F)<sup>a,b</sup> | *n* | Maximum adult length and mass<sup>b</sup> |
| --- | --- | --- | --- | --- |
| *Eubalaena australis* | Southern Right Whale | 14.3 m / 14.8 m<br>46.7 t / 52.0 t | 1,341<sup>c</sup> / 551 | 17.4 m<br>85.9 t |
| *Eubalaena japonica* | North Pacific Right Whale | 16.0 m / 16.8 m<br>66.2 t / 77.1 t | 71 / 40 | 19.7 m<br>126.4 t |
| *Balaena mysticetus* | Bowhead Whale | 14.4 m / 15.4 m<br>47.7 t / 58.8 t | 200<sup>c</sup> / 234<sup>c</sup> | 20.5 m<br>172.0 t |
| *Eschrichtius robustus* | Gray Whale | 12.0 m / 12.6 m<br>19.1 t / 21.9 t | 792<sup>c</sup> / 773<sup>c</sup> | 15.2 m<br>39.3 t |
| *Megaptera novaeangliae* | Humpback Whale | 12.4 m / 13.3 m<br>27.7 t / 34.1 t | 31,324<sup>c</sup> / 7,237 | 18.0 m<br>83.7 t |
| *Balaenoptera edeni* | Bryde's Whale | 12.6 m / 13.1 m<br>14.1 t / 15.7 t | 6,594<sup>c</sup> / 2,177 | 16.3 m<br>28.6 t |
| *Balaenoptera borealis* | Sei Whale | 14.4 m / 15.4 m<br>23.1 t / 28.6 t | 85,055<sup>c</sup> / 39,843 | 20.1 m<br>66.5 t |
| *Balaenoptera physalus* | Fin Whale | 20.1 m / 21.7 m | 204,370<sup>c</sup> / 99,863 | 27.2 m |
|  |  | 51.9 t / 63.0 t |  | 112.1 t |
| *Balaenoptera musculus* | Blue Whale | 23.6 m / 25.1 m | 34,129<sup>c</sup> / 12,616 | 31.1 m |
|  |  | 84.4 t / 103.3 t |  | 209.0 t |

- a. See text and subsequent tables as some mean lengths are weighted averages for populations with significant size differences. - b. Body masses are estimated from either linear regression or largest oil yields. - c. Approximate sample size from component curves

Most of the published mass data for large mysticetes were the sums of individual pieces weighed on platform scales or bulk fillings of pressure cookers (Lockyer, 1976). Blood and other body fluids are lost in this process, typically estimated to compose 6-7% of the total body mass (Laurie, 1933; Moore et al., 2004). The canonical allometric formula (Equations 1 & 2) and the Rice-Wolman (RW) model (Equation 3) are reported for mass relationships (George, 2009; Rice & Wolman, 1971; Schultz, 1938). Formulas, data, references, and code are provided in Supplementary Files 1-4, respectively.

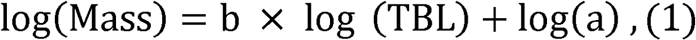

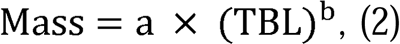

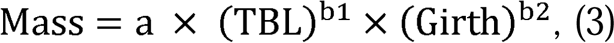

As in the analysis for odontocetes (McClure, 2024), the mean adult TBL (*L*_mean_) and standard deviation (SD) of females were primarily estimated using direct samples of individuals recorded as sexually mature in the commercial data. Corresponding parameters for males were estimated by dissecting length frequencies of mature individuals from normal probability plots (Harding, 1949). To enforce objectivity and consistency, published estimates for the *L*_sm_ were used as starting values for the point of inflection. Preliminary comparisons for select female catches validated the congruency between the two methods. Therefore, the graphical analysis was applied to certain female catches where the difference in sample size deemed it preferrable. Regional differences in the *L*_mean_ values were examined using two-sample *t*-tests (Zar, 2010).

In general, the regional differences in the *L*_mean_ for most rorqual species, even when statistically significant (p < 0.05), were consistent with previous observations of notable morphological differences (Best, 1977; Best & Lockyer, 2002; Lockyer, 1977) or often relatively minor regional or temporal variation in asymptotic size and growth rate (Lockyer, 1981; Mogoe et al., 2014; Ohsumi, 1980; Panfilov, 1978). However, differences were also detected for populations that were considered synonymous in size, such as within *M. novaeangliae* (Chittleborough, 1965; Stevick, 1999). Given limitations of such comparisons against confounding factors such as precise age composition (McClure, 2024), the results section will mainly highlight more convincing signs of morphological differences or similarities not illustrated in previous research.

The analyses for rorqual species within in the IWC catch database are restricted to expeditions from after the introduction of minimum length regulations in 1937, corresponding to the ‘’Late period’’ as described in previous work on *B. musculus* (Branch et al., 2007). This convention helps minimize data quality issues within early whaling data such as non-standard measuring techniques, visual estimation, and rounding to 5-ft intervals (Branch et al., 2007; Donovan, 2000). While the intentional ‘stretching’ of whales below the minimum legal size is known to afflict much of the data reported prior to the IOS (Best, 1989), these size restrictions had minimal overlap with the distributions of mature individuals.

The current review distinguishes the three species within *Eubalaena* (Rosenbaum et al., 2000): *E. glacialis* in the North Atlantic (Müller, 1776), *E. japonica* in the North Pacific (Lacépède, 1818) and *E. australis* in the Southern Hemisphere (Desmoulins, 1822). Due to limited samples across both modern catches and necropsies, the adult size for *E. glacilias* was not as deeply examined. The *L*_mean_ for *E. australis* and *B. mysticetus* were respectively calculated from the Soviet commercial catches (Figure S1) and North Pacific subsistence harvests recorded in the IWC catch database (Figure S2). The *L*_mean_ for *E. japonica* (Table 2 & Figure S1) was calculated from data on sexed individuals from literature (Ivashchenko et al., 2017; Ivashchenko & Clapham, 2012; Klumov, 1962; Omura et al., 1969; Pike & MacAskie, 1969; Zenkovich, 1954). All non-examined specimens ≥ 15 m were assumed to be sexually mature. Midpoints were used for lengths binned to 50 cm intervals.

**Table 2:** Growth parameters for Balaenidae.

| Species | $L_b$ (m) | $L_w$ (m) | $L_{sm}$ (m)<br>M/F | $L_{mean} \pm SD$ (m)<br>M/F | $L_{\infty}$ (m)<br>M/F | $L_{max}$ (m)<br>M/F | Reference |
| --- | --- | --- | --- | --- | --- | --- | --- |
| *E. japonica* | — | 11 | 14.5<sup>a</sup> | 16.0 ± 0.86 /<br>16.8 ± 1.13 | 16.5<sup>a</sup> / 17.5<sup>a</sup> | 18.3<sup>a</sup> / 19.7<sup>a</sup> | Ivashchenko & Clapham, 2012; Klumov, 1962; Omura et al., 1969; Current study |
| *E. australis* | 4.7 | 9.5 | 12.6<sup>b</sup> / 13.1 | 14.3 ± 0.78<sup>c</sup> /<br>14.8 ± 0.98<sup>c</sup> | 13.7<sup>b</sup> / 14.3 | 17.2 / 17.4 | Christiansen et al., 2022; Current study |
| *B. mysticetus* | 4.3 | 8.5 | 12.7<sup>a</sup> / 13.4 | 14.4 ± 1.20 /<br>15.4 ± 1.26<sup>d</sup> | 15.5<sup>a</sup> / 16.5<sup>a</sup> | 19.2<sup>e</sup> / 20.5<sup>e</sup> | George et al., 1999; Lubetkin et al., 2012; Tarpley et al., 2021; Current study |

- a. Midpoint of binned value or estimated range in literature. - b. Estimated by adjusting for 4% difference between males and females - c. Whaling data may not be directly comparable to $L_{\infty}$ values from photogrammetry surveys. - d. Estimated from graphical analysis of total female catch. - e. Estimates from Equation 4.

All stocks within the *B. edeni* complex are treated as one species as the precise taxonomy has yet to be resolved (Constantine et al., 2018; Junge, 1950). Due to low sample sizes, *B. edeni* catches from the South Pacific, Southern land stations, and Indian Ocean were excluded from analyses. Distinct subspecies corresponding to the Northern and Southern Hemispheres are recognized for nearly all other rorqual species (Jackson et al., 2014; Rice, 1998; Rosel & Wilcox, 2014). Throughout lower latitudes, the southern populations of *B. musculus* are delineated into *B. m. brevicauda* (Ichihara, 1966), *B. m. indica* (Blyth, 1859), and another proposed subspecies in Southeast (SE) Pacific (Attard et al., 2024; Pastene et al., 2020). Each are distinct in size from *B. m. intermedia* (Burmeister, 1871), the primary subspecies of the Antarctic.

Since mixture within catches would confound regional comparisons, the *L*_mean_ values for *B. musculus* in the Southern Hemisphere is summarized in accordance with subspecies rather than ocean basin. Parameters for mature females from the SE Pacific and the Antarctic were obtained directly from a previous analysis (Branch et al., 2007). For the graphical analysis of Antarctic males, whales caught below 60**°**S were isolated to minimize overlap with other subspecies. Analyses for *B. m. brevicauda* were restricted to data from Soviet fleets that ignored the 21.3 m minimum length restriction. Due to known morphological similarities (Branch & Mikhalev, 2008), *B. m. indica* is synonymized with *B. m. brevicauda* in this review.

Since the length distributions for mature mysticetes are approximately normal (Branch et al., 2007; Ohsumi et al., 1958), the expected upper bound for the maximum value within a normally-distributed sample of *n* (Equation 4) is used as an approximation to compare against potentially unreliable maximum values within the whaling data (Kamath, 2015). For a given species/subspecies, *n* is the total number of whales in the entire IWC catch database that were greater than or equal to the *L*_sm_ and rounded to the nearest thousand (*Y* = deviation from mean TBL).

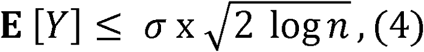

Since natural variation may enable the occurrence of unexpected outliers, reliable estimates for the *L*_max_ will also be determined using a combination of the verified measurements in literature and the largest sizes recorded across different expeditions of the total catches within the commercial data. This helps to account for the possible influence of biases in measuring technique.

## 3 Review of size distributions and mass relationships

### Balaenidae

#### Total Body Length

Despite similarity in size between *E. glacilias* and *E. australis* up to sexual maturity (Christiansen et al., 2022; Fortune et al., 2021), mature females are known to be significantly larger and in better body condition in the Southern Hemisphere due to environmental stressors (Christiansen et al., 2020; Stewart et al., 2021). Aside from *E. japonica* being significantly larger, differences in external morphology between the three species are minimal (Fortune et al., 2021; Omura et al., 1969). Table 2 shows that the *L*_mean_ for *E. australis* from the whaling dataset exceeded the photogrammetric estimates for the *L*^∞^ (Christiansen et al., 2022). This appears to reflect previous concerns regarding the compatibility between photogrammetric and whaling data for this species (Tormosov et al., 1998).

*E. glacialis* is sometimes cited to reach a maximum TBL of 17.8-18 m (Cummings, 1985; Huang et al., 2009), but these measurements were either unstandardized or taken from *E. japonica* when the two species were still synonymized (Omura, 1958; Thompson, 1928). The largest reliable measurement for *E. glacialis* was a 16.5 m female captured near Long Island, New York (Andrews, 1908). The Soviet data for *E. australis* reported males reaching 17.2 m, and the infraction records for other nations reported a female measuring 17.4 m. The largest reliable measurements for *E. japonica* appear to be a binned length of 18.1-18.5 m for a male and 19.5-19.8 m for a female (Ivashchenko & Clapham, 2012).

For *B. mysticetus*, the largest female and male within the analyzed harvest data were 19.2 m and 17.8 m, respectively. The minimum estimate for the historical population size of the combined Bering-Chukchi-Beaufort and Okhotsk stocks of *B. mysticetus* is around 13,000 (Woodby & Botkin, 1993). If it’s assumed that mature females composed 25% of this value, Equation 4 predicts the female *L*_max_ to be 20.5 m. Given the selection against larger individuals in subsistence harvests and the high variability of continued growth in *B. mysticetus*, Equation 4’s prediction is likely the best *L*_max_ value for this species.

#### Total Body Mass

Postnatal and near-term fetal weights have been published for *E. japonica* (n= 18), *E. glacialis* (n=13), *E australis* (n=3), and *B. mysticetus* (n=7) from necropsies and whaling (Best, 2008; Fortune et al., 2021; George, 2009; Klumov, 1962; Omura et al., 1969). The heaviest record for *E. japonica* was a 17.4 m lactating female that weighed 106.5 tonnes after flensing or 113.3 tonnes after a 6 % fluid correction (Klumov, 1962). The heaviest RW body mass estimate for *B. mysticetus* was 101 tonnes for a 16.5 m female (George et al., 2021).

The RW1 body mass model published for *B. mysticetus* (George, 2009) accurately predicted the body masses for the non-pregnant *E. japonica* specimens (r^2^= 0.969; n=6), excluding three outliers with known errors in girth measurements (Christiansen et al., 2019). AIC model selection preferred a linear model (Equation 1) that pooled the mass data for the two genera (AICc = -65.98, weight = 0.74). Mass data and external measurements for each species were pooled into Equations 1-3 as composite models for the Balaenidae family (Table S1, Figures 1 & S3). As explicitly noted by the original author (Klumov, 1962), corrections on the order of 8-11 tonnes were necessary for three whales that were missing components for the skulls and viscera. This went unmentioned when the data was republished in the more widely-cited review (Omura et al., 1969) and uncorrected in subsequent literature (Fortune et al., 2021; Lockyer, 1976).

**Figure 1:**
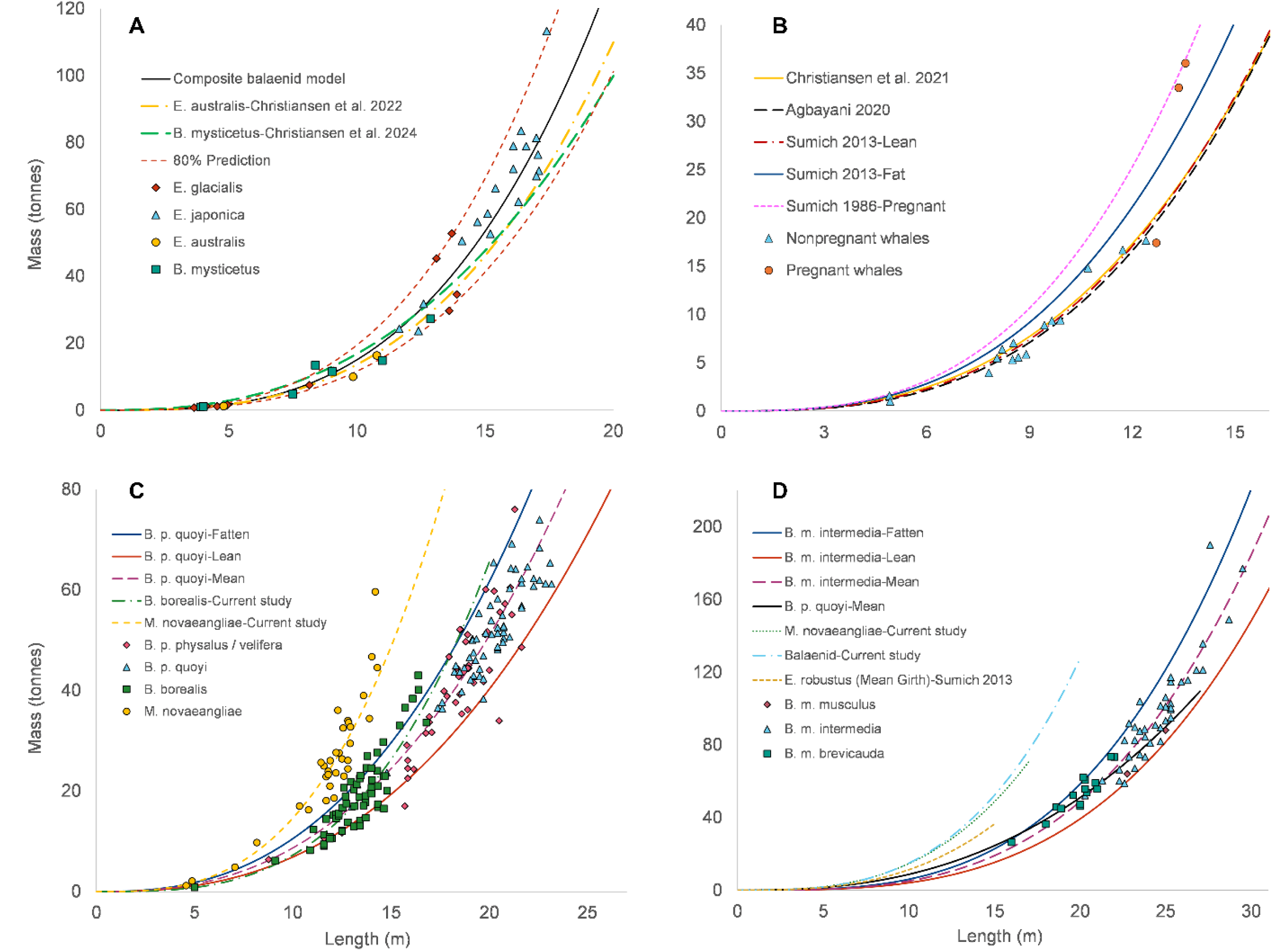
Body mass relationships for large mysticetes. Legend: Fluid-corrected body mass data for Balaenidae (A), *E. robustus* (B), *M*. *novaeangliae*/*B. physalus/B. borealis* (C), and *B. musculus* (D).

Figure 1A compares the composite balaenid model (Equation 2) to photogrammetric mass models for *E. australis* and *B. mysticetus* (Christiansen et al., 2022, 2024). A balaenid would weigh 54.2 tonnes (80% PI 42.6-69.4 tonnes) at 15 m and 143.0 tonnes (80% PI 111.1-184.2 tonnes) at 20.5 m. Oil yield to body mass conversions for *B. mysticetus* (George et al., 2007, 2021) suggest that individual yields 337-375 barrels (37.0-40.9 tonnes) of oil (Fairburn, 1955; Pease, 1918; Williams, 1904) corresponded to intact body masses of 154-172 tonnes. These figures fall well within the expected body mass for a 20.5 m balaenid.

### Eschrichtiidae

#### Total Body Length

Table 3 summarizes the growth parameters for the ENP population of *E. robustus* (Agbayani et al., 2020), which is thought to be synonymous in size to the WNP population. The physically smaller Pacific Coast Feeding Group (PCFG) subpopulation (Bierlich et al., 2023) are excluded due to limited historical catch data. Graphical analysis of the total catches was employed for both sexes (Table 3 & Figure S4). The largest canonically cited individuals are a 14.62 m male and a 15.54 m unsexed specimen (Cope, 1868), though the reliability of these measurements are questionable (Andrews, 1914). A 14.3 m male and 15 m female were verified by researchers aboard the *Aleut* (Zenkovich, 1934, 1937). The commercial dataset reports 10 males exceeding 14.3 m and females reaching up to 15.2 m. When accounting for sexual dimorphism, the largest reliably recorded males in the commercial data appear to be 14.5 m.

**Table 3:**
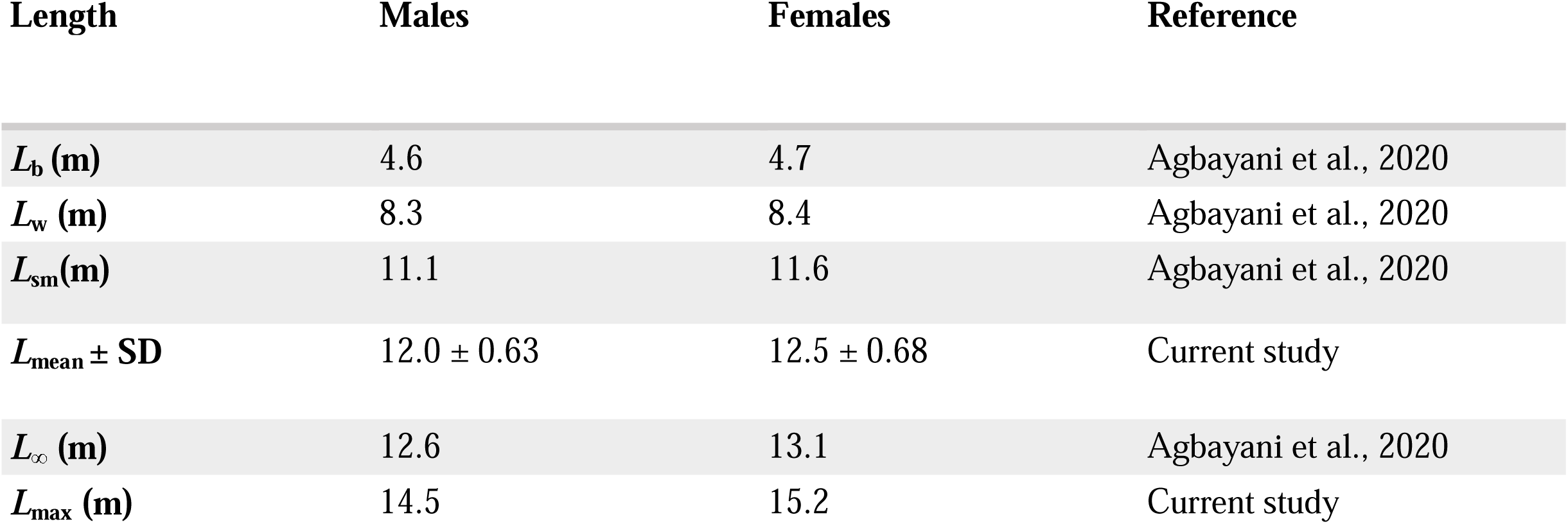
Growth parameters for *E. robustus*.

| Length | Males | Females | Reference |
| --- | --- | --- | --- |
| $L_b$ (m) | 4.6 | 4.7 | Agbayani et al., 2020 |
| $L_w$ (m) | 8.3 | 8.4 | Agbayani et al., 2020 |
| $L_{sm}$ (m) | 11.1 | 11.6 | Agbayani et al., 2020 |
| $L_{mean} \pm SD$ | $12.0 \pm 0.63$ | $12.5 \pm 0.68$ | Current study |
| $L_{\infty}$ (m) | 12.6 | 13.1 | Agbayani et al., 2020 |
| $L_{max}$ (m) | 14.5 | 15.2 | Current study |

#### Total Body Mass

In total, 36 body masses for *E. robustus* collected from whaling, strandings, and captive specimens have been published (Agbayani et al., 2020; Sumich et al., 2013). The heaviest was a 13.55 m pregnant female that weighed 33.85 tonnes after flensing (Rice & Wolman, 1971). The various published models for this species are provided in Table S2 (Agbayani et al., 2020; Christiansen et al., 2021; Lockyer, 1976; Sumich, 1986; Sumich et al., 2013). For the RW curves in Figure 1B, girths are fixed to the observed values of 50% of the TBL for the lean condition, 60% for the standard fattened condition, and 70% for the pregnant fattened condition (Sumich, 1986).

Figure 1B shows that the mass relationship published alongside a recent growth curve (Agbayani et al., 2020) precisely aligned with both the RW curve for the lean condition and a volume model for a predominantly emaciated sample of whales (Christiansen et al., 2021). This supports concerns by the authors (Agbayani et al., 2020) that the sample for this model appears too limited to represent the normal condition. When setting the RW model at the normal body condition (girth = 55% TBL), *E. robustus* would weigh 19.1 tonnes at 12 m while a pregnant female would weigh 39.3 tonnes at 15.2 m.

### Balaenopteridae

#### Total Body Length

Growth parameters along with corresponding *L*_mean_ values are provided in Table 4. Due to congruity in existing *L*_sm_ and *L*^∞^ estimates, all *M. novaeangliae* populations and certain subspecies within *Balaenoptera* were pooled together in Table 4 and supplementary plots (Figures S5-S17).

**Table 4:**
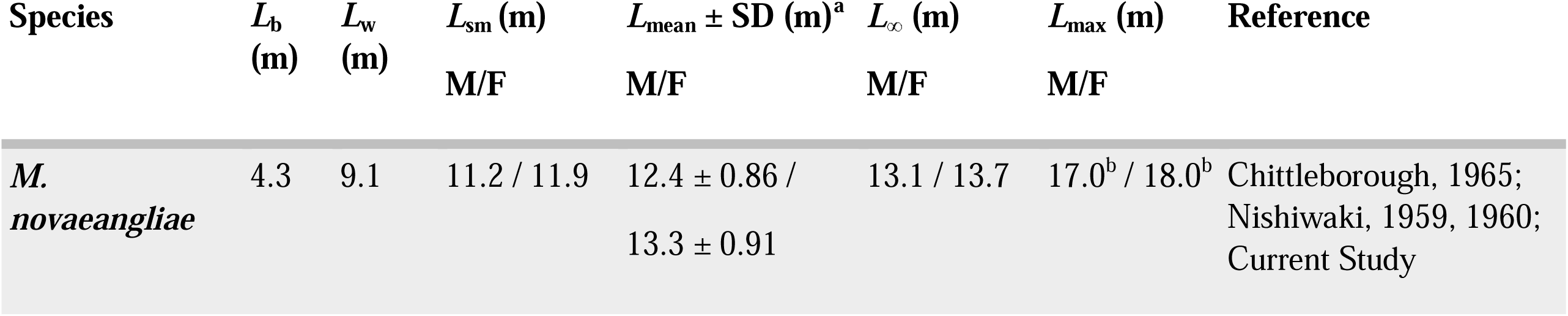
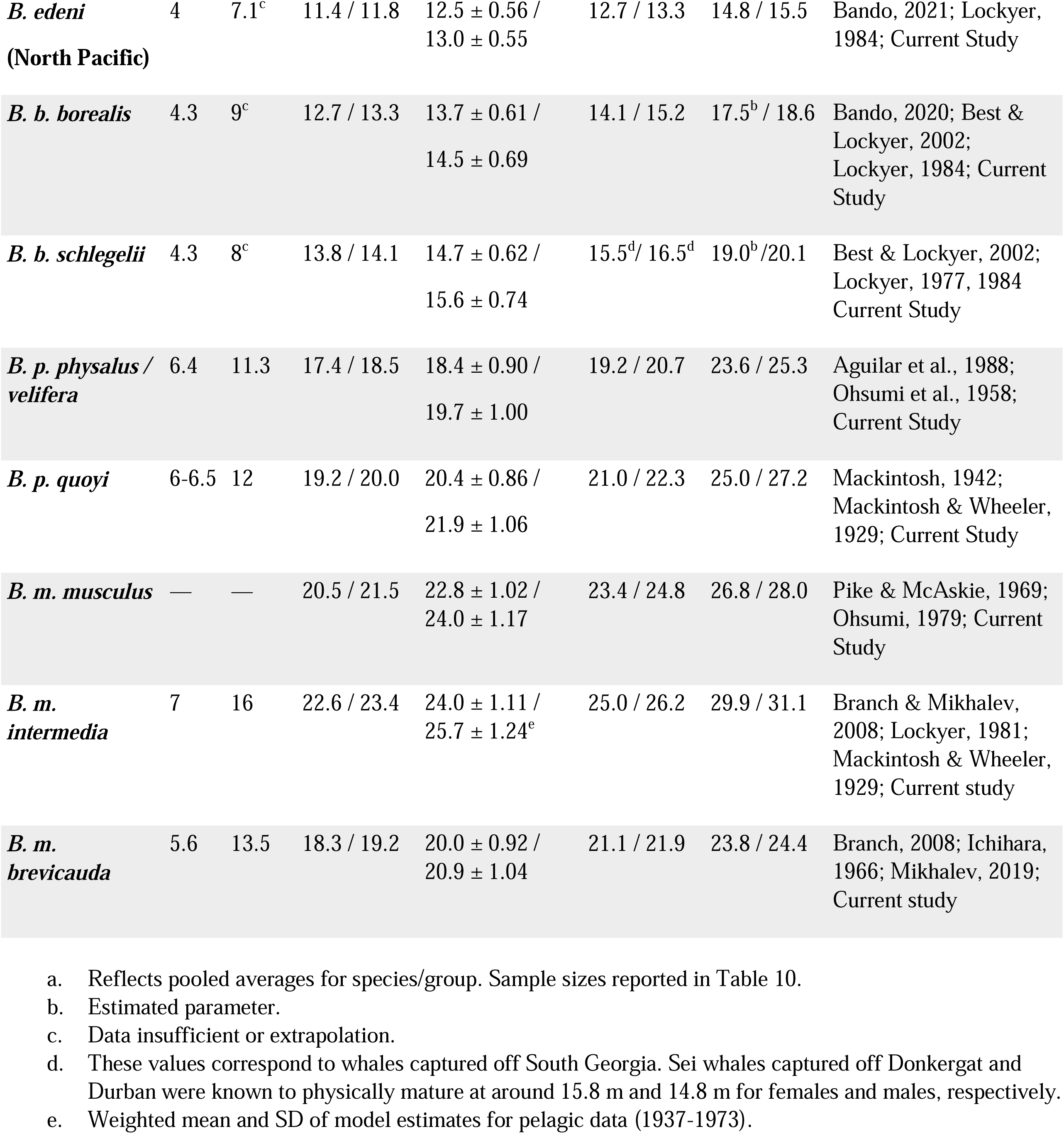
Growth parameters for Balaenopteridae.

| Species | $L_b$<br>(m) | $L_w$<br>(m) | $L_{\text{sm}}$ (m)<br>M/F | $L_{\text{mean}} \pm \text{SD}$ (m)<sup>a</sup><br>M/F | $L_{\infty}$ (m)<br>M/F | $L_{\text{max}}$ (m)<br>M/F | Reference |
| --- | --- | --- | --- | --- | --- | --- | --- |
| *M. novaeangliae* | 4.3 | 9.1 | 11.2 / 11.9 | 12.4 ± 0.86 /<br>13.3 ± 0.91 | 13.1 / 13.7 | 17.0<sup>b</sup> / 18.0<sup>b</sup> | Chittleborough, 1965;<br>Nishiwaki, 1959, 1960;<br>Current Study |
| *B. edeni*<br>(North Pacific) | 4 | 7.1<sup>c</sup> | 11.4 / 11.8 | 12.5 ± 0.56 /<br>13.0 ± 0.55 | 12.7 / 13.3 | 14.8 / 15.5 | Bando, 2021; Lockyer,<br>1984; Current Study |
| *B. b. borealis* | 4.3 | 9<sup>c</sup> | 12.7 / 13.3 | 13.7 ± 0.61 /<br>14.5 ± 0.69 | 14.1 / 15.2 | 17.5<sup>b</sup> / 18.6 | Bando, 2020; Best &<br>Lockyer, 2002;<br>Lockyer, 1984; Current<br>Study |
| *B. b. schlegelii* | 4.3 | 8<sup>c</sup> | 13.8 / 14.1 | 14.7 ± 0.62 /<br>15.6 ± 0.74 | 15.5<sup>d</sup> / 16.5<sup>d</sup> | 19.0<sup>b</sup> / 20.1 | Best & Lockyer, 2002;<br>Lockyer, 1977, 1984<br>Current Study |
| *B. p. physalus* /<br>*velifera* | 6.4 | 11.3 | 17.4 / 18.5 | 18.4 ± 0.90 /<br>19.7 ± 1.00 | 19.2 / 20.7 | 23.6 / 25.3 | Aguilar et al., 1988;<br>Ohsumi et al., 1958;<br>Current Study |
| *B. p. quoyi* | 6-6.5 | 12 | 19.2 / 20.0 | 20.4 ± 0.86 /<br>21.9 ± 1.06 | 21.0 / 22.3 | 25.0 / 27.2 | Mackintosh, 1942;<br>Mackintosh & Wheeler,<br>1929; Current Study |
| *B. m. musculus* | — | — | 20.5 / 21.5 | 22.8 ± 1.02 /<br>24.0 ± 1.17 | 23.4 / 24.8 | 26.8 / 28.0 | Pike & McAskie, 1969;<br>Ohsumi, 1979; Current<br>Study |
| *B. m.*<br>*intermedia* | 7 | 16 | 22.6 / 23.4 | 24.0 ± 1.11 /<br>25.7 ± 1.24<sup>e</sup> | 25.0 / 26.2 | 29.9 / 31.1 | Branch & Mikhalev,<br>2008; Lockyer, 1981;<br>Mackintosh & Wheeler,<br>1929; Current study |
| *B. m.*<br>*brevicauda* | 5.6 | 13.5 | 18.3 / 19.2 | 20.0 ± 0.92 /<br>20.9 ± 1.04 | 21.1 / 21.9 | 23.8 / 24.4 | Branch, 2008; Ichihara,<br>1966; Mikhalev, 2019;<br>Current study |

For *M. novaeangliae*, Equation 4 predicted *L*_max_ values of 17.5 m (n = 34,000) for females and 16.4 m (n = 44,000) for males. Since 1937, 17 males and 13 females were reported to measure 17 m or greater. The largest female was reportedly 19.5 m, while the next 13 were 18.3 m or lower.

The largest record amongst precisely calibrated photogrammetry measurements was a 16.8 m individual observed repeatedly in the Arabian Sea (F. Christiansen, personal communication, 2023). The mounted skeleton displayed at the Nha Trang Institute of Oceanography in Vietnam appears to be the largest in the world, officially measuring 18 m (Redman, 2019). When comparing this to the values from the commercial data and Equation 4’s predictions, the *L*_max_ for *M. novaeangliae* can be estimated to be 18 m for females and 17 m for males.

**Table 5:** Regional comparison of *L*_mean_ for *M. novaeangliae*.

| Region | $L_{\text{mean}} \pm \text{SD (m)}$ | |
| --- | --- | --- |
|  | Males | Females |
| North Pacific | 12.4 ± 0.93<br>(2,594) | 13.2 ± 0.97<br>(465) |
| North Atlantic | 12.9 ± 1.16<br>(113) | 13.7 ± 0.90<br>(35) |
| South Pacific | 12.4 ± 0.70<br>(6,008) | 13.3 ± 0.74<br>(433) |
| South Atlantic | 13.1 ± 1.00<br>(1,805) | 14.4 ± 0.93<br>(62) |
| Indian Ocean | 12.2 ± 0.79<br>(8,756) | 13.3 ± 0.80<br>(632) |
| S. Hemisphere Land Stations | 12.8 ± 0.78<br>(178) | 13.8 ± 0.91<br>(61) |
| S. Hemisphere Pelagic | 12.4 ± 0.78<br>(5,781) | 13.2 ± 0.87<br>(3,455) |
| S. Hemisphere Pelagic<br>(Soviet) | 12.4 ± 1.00<br>(4,740) | 13.3 ± 0.96<br>(2,097) |

For *B. edeni*, the length distributions of mature females in the South Atlantic and Southern pelagic catches were polymodal (Figures S6 & S7), consistent with observations of distinct morphs (Best, 1977; Ohsumi, 1980). The South Atlantic data contains a peak around 15.5 m and a maximum TBL of 18 m (Figure S6), indicating that nearly 20% of the sample were misidentified catches of *B. borealis* (Figure S7). Outliers of this type comprised only a small percentage (< 2.0 %) of the total male catches for the South Atlantic and North Pacific. Otherwise, no obvious mixtures of different morphs or species were detected in other regions (Figures S8 & S9).

Due to rampant misidentification with other rorquals, caution should be taken with commercial records of individuals exceeding 15.5 m, a length typically not exceeded in verified records (Best, 1977; Ohsumi, 1980). The largest specimen that can be tentatively assigned to *B. edeni* was a 16.3 m female captured off of Western Australia in 1963 (Best, 1977). A corresponding record of a 15.9 m male within the Soviet catch data from the Gulf of Aden seemed particularly convincing, as both the whaling data and lack of confirmed sightings in literature suggests that sei whales are infrequent or wholly absent in the Arabian Sea region.

**Table 6:**
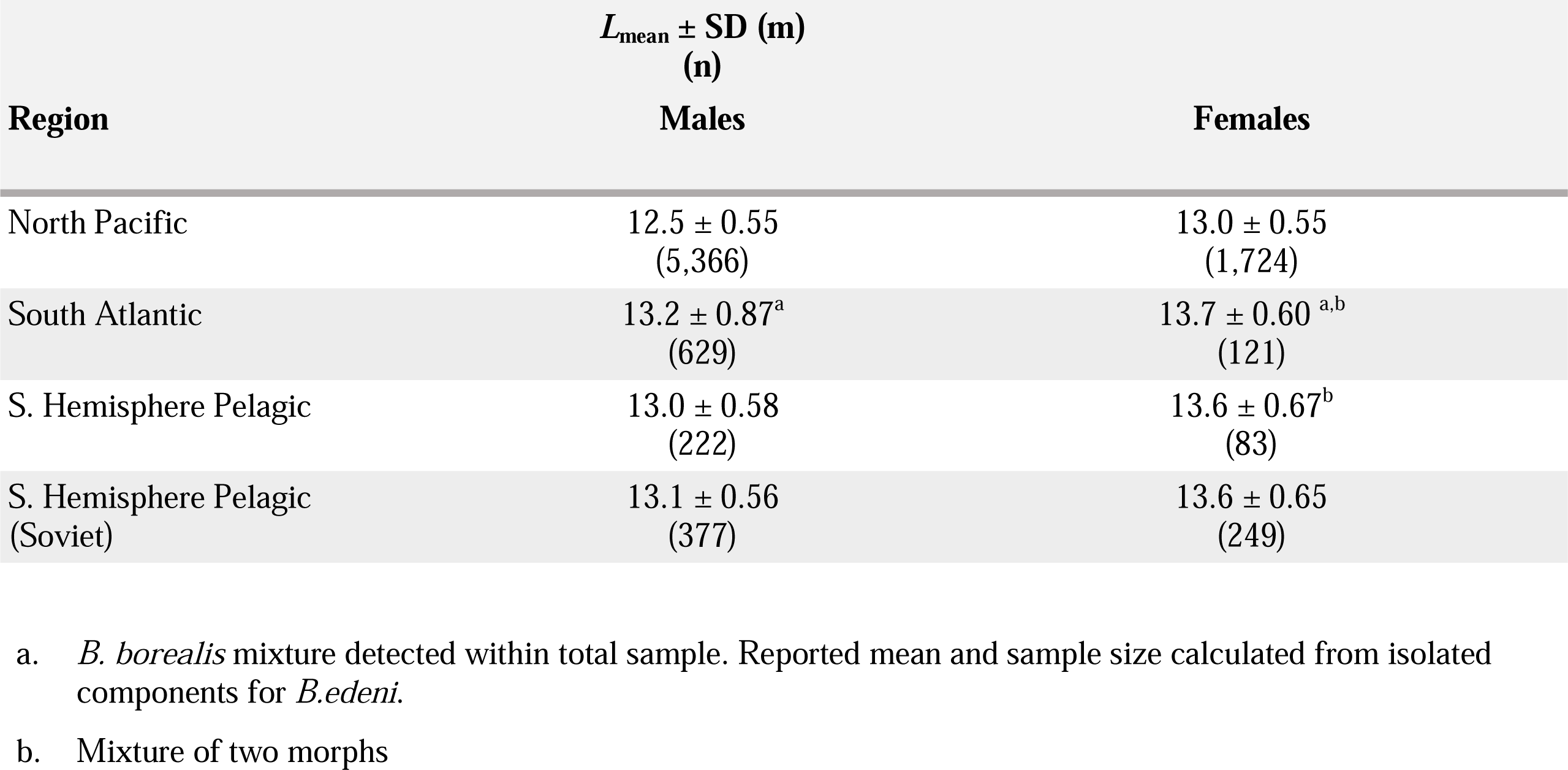
Regional comparison of *L*_mean_ for *B. edeni*.

| Region | $L_{\text{mean}} \pm \text{SD (m)}$<br>(n) | |
| --- | --- | --- |
|  | Males | Females |
| North Pacific | 12.5 ± 0.55<br>(5,366) | 13.0 ± 0.55<br>(1,724) |
| South Atlantic | 13.2 ± 0.87<sup>a</sup><br>(629) | 13.7 ± 0.60<sup>a,b</sup><br>(121) |
| S. Hemisphere Pelagic | 13.0 ± 0.58<br>(222) | 13.6 ± 0.67<sup>b</sup><br>(83) |
| S. Hemisphere Pelagic<br>(Soviet) | 13.1 ± 0.56<br>(377) | 13.6 ± 0.65<br>(249) |

a. *B. borealis* mixture detected within total sample. Reported mean and sample size calculated from isolated components for *B. edeni*.
b. Mixture of two morphs

The *L*_mean_ values for *B. borealis* from the South Pacific were lowest in the Southern Hemisphere, falling within the size range of adults from the Northern Hemisphere. The graphically estimated parameters reported in Table 7 were consistent with the limited sample of mature females (mean = 14.1 m; SD = 0.76 m; median & mode=14.3 m; *n* = 26). The data from the South Pacific was not used to calculate the *L*_mean_ for southern sei whales in Table 4.

**Table 7:** Regional comparison of *L*_mean_ for *B. borealis*.

| Region | $L_{\text{mean}} \pm \text{SD (m)}$ | |
| --- | --- | --- |
|  | Males | Females |
| North Pacific | 13.7 ± 0.61<br>(18,588) | 14.6 ± 0.68<br>(6,850) |
| North Atlantic | 13.3 ± 0.65 | 14.2 ± 0.65 |

Body size of large mysticetes
|  | (1,783) | (315) |
| --- | --- | --- |
| South Pacific | 13.6 ± 0.90<br>(284) | 14.4 ± 0.80<sup>a</sup><br>(362) |
| South Atlantic | 14.6 ± 0.90<br>(3,892) | 15.0 ± 0.85<br>(1,145) |
| Indian Ocean | 14.0 ± 0.68<br>(1,212) | 15.0 ± 0.94<br>(102) |
| S. Hemisphere Land Stations | 14.9 ± 0.55<br>(4,825) | 15.8 ± 0.68<br>(2,692) |
| S. Hemisphere Pelagic | 14.6 ± 0.59<br>(42,127) | 15.6 ± 0.73<br>(20,330) |
| S. Hemisphere Pelagic<br>(Soviet) | 14.8 ± 0.65<br>(13,393) | 15.6 ± 0.76<br>(8,407) |

a. Estimated from graphical analysis of total female catch

The largest female within the total catches for *B. borealis* in the Northern Hemisphere was an 18.6 m individual captured off the North Pacific. The next 13 measured between 17-18 m. The largest in the Southern Hemisphere were two 20.1 m individuals captured at South Georgia in 1949 and 1960. The next 15 largest measured between 18.3 m-19.6 m. These do not readily appear to be misidentified records for *B. physalus* when adding some allowance to the Equation 4 predictions of 17.6 m (n = 18,000) for northern females and 19.1 m (n = 67,000) for southern females. There are questionable records of males exceeding the maximum reported female size. The *L*_max_ for males reported in Table 4 were therefore downsized from the largest females using the ratio of sexual dimorphism of their respective *L*^∞^ values.

Despite possessing similar means (Table 8), the female length distributions for *B. physalus* in the South Atlantic and Indian Ocean land stations were strongly negatively skewed (Figure S13). While the distributions for the rest of the Southern Hemisphere were approximately normal, outliers as small as 14.3-16 m were detected across the Chilean, Soviet, and British catches.

**Table 8:** Regional comparison of *L*_mean_ for *B. physalus*.

| Region | $L_{\text{mean}} \pm \text{SD (m)}$ | |
| --- | --- | --- |
|  | Males | Females |
| North Pacific | 18.4 ± 0.89<br>(16,619) | 19.7 ± 0.94<br>(4,877) |
| North Atlantic | 18.2 ± 1.01<br>(12,593) | 19.6 ± 1.10<br>(2,446) |
| South Pacific | 20.1 ± 0.78<br>(142) | 21.3 ± 1.00<sup>a</sup><br>(94) |
| South Atlantic | 19.9 ± 1.19<br>(666) | 21.3 ± 2.07<sup>b</sup><br>(35) |
| Indian Ocean | 19.9 ± 0.99<br>(3,169) | 21.8 ± 1.47<sup>b</sup><br>(73) |
| S. Hemisphere Land Stations | 19.9 ± 0.91<br>(14,564) | 21.7 ± 1.14<br>(3,892) |
| S. Hemisphere Pelagic | 20.4 ± 0.82<br>(173,095) | 21.9 ± 1.05<br>(85,403) |
| S. Hemisphere Pelagic<br>(Soviet) | 20.1 ± 1.00<br>(4,828) | 21.5 ± 1.19<br>(3,214) |

a. Estimated from graphical analysis of total female catch
b. Negatively skewed.

The Equation 4 predictions for northern and southern females were respectively 24.2 m (n = 29,000) and 27.2 m (n = 220,000). One photogrammetry survey of the North Atlantic subspecies verified an unsexed individual measuring 24.45 m (Mészáros et al., 2025), with the whaling dataset reporting a female measuring 25 m. Another female from the North Pacific whaling dataset measured 25.3 m. The largest male from the North Pacific measured 23.6 m, congruent with the ratio of sexual dimorphism between the *L*_mean_ values in Table 8. The Soviet data from the Antarctic records one female reaching 27.2 m, consistent with Equation 4’s prediction. A southern male reported to measure 25 m appears to be the largest record that’s congruent with the observed sexual dimorphism.

The SE Pacific subspecies of *B. musculus* was very comparable in size to the northern subspecies (Table 9). However, a significant difference was detected between the SE Pacific and North Pacific (*t-*test, *p* < 0.05). The largest whales reliably identified as *B. m. brevicauda* appear to have been around 24.2-24.5 m (Omura, 1984; Pastene et al., 2020). The largest individual in the Northern Hemisphere dataset was a 28.0 m female from the North Atlantic, a size substantiated by mounted skeletons and photogrammetry studies on the northern subspecies (Gorter, 2010; Ortega-Ortiz et al., 2022).

**Table 9:** Regional comparison of *L*_mean_ for *B. musculus*.

| Region | $L_{\text{mean}} \pm \text{SD (m)}$ | |
| --- | --- | --- |
|  | Males | Females |
| North Pacific | $22.8 \pm 0.98$<br>(1,054) | $24.1 \pm 1.05$<br>(183) |
| North Atlantic | $22.2 \pm 1.03$<br>(202) | $23.4 \pm 1.45$<br>(44) |
| Antarctic | $24.0 \pm 1.11$<br>(28,865) | $25.7 \pm 1.24^a$<br>(10,809) |
| Pygmy | $20.0 \pm 0.92$<br>(3,199) | $20.9 \pm 1.04$<br>(1,509) |
| SE Pacific | $22.1 \pm 1.13^b$<br>(809) | $23.5 \pm 1.20^b$<br>(71) |

a. Weighted average of mean and SD of Pelagic catch below 60°S (1937-1973) from Branch et al., 2007.
b. Includes data taken since 1931.

While the literature (Risting, 1922, 1928) and commercial data from early Norwegian expeditions report that *B. m. intermedia* reached 32-35 m. The maximum sizes within Norwegian catch statistics fell in line with other nations after 1931, hinting that at least most of these exceptional records were artefacts from measuring procedures. The greatest TBL for *B. m. intermedia* to be directly measured by a scientist was 29.9 m (Rice, 1978). Since 1937, the commercial data reports six females ≥ 30.5 m, the largest was taken in 1948 and measured 31.1 m. This size falls well within Equation 4’s prediction of 31.6 m when conservatively estimating that only 50% of all captured females in the IWC catch database were mature (n = 77,956).

#### Total Body Mass

The heaviest summed piecemeal masses reported in communications and previous literature were 56.1 tonnes for *M. novaeangliae* (Best, 2008), 24.9 tonnes for *B. edeni* (Tamura et al., 2016), and 40.45 tonnes for *B. borealis* (Y. Ivashchenko, personal communication, 2023). The heaviest intact body mass for *B. physalus* was a 21.3 m North Atlantic female that weighed 76.3 tonnes with corrections for missing tongue and body fluids (Víkingsson et al., 1988). The heaviest record for *B. musculus* was a 27.6 m pregnant female that weighed 190 tonnes (Voronin, 1948). This individual’s maximum blubber thickness of 42 cm substantiates an exceptional body condition against hundreds of other whales (Slijper, 1948). A record of 196 long tons / 199 tonnes reported in some literature (Carwardine, 1995; McClain et al., 2015), appears to be a conversion error for a total mass of 178 tonnes /196 short tons (Scheffer, 1974).

Along with existing formulas (Table S8), body mass data for *M. novaeangliae* (n = 50), *B. borealis* (n = 60), *B. physalus* (n =106) and *B. musculus* (n = 56) were compiled for this review. Fluid-corrected body mass data and corresponding curves for Balaenopteridae are compared in Figures 1C & D. The dataset for *M. novaeangliae* and *B. borealis* were larger than those available to previous authors (Lockyer, 1976; Mikhalev, 2019) and all data were compiled into mass models for their respective species (Table S3, Figures 1, S18 & S19). While AIC model selection strongly preferred separate regional models for *B. borealis* (AICc=-145.32, weight = 1), summary statistics from unpublished JARPNII survey data (Tamura et al., 2016) strongly hint that previously observed regional differences in body mass were due to temporal sampling bias (Víkingsson et al., 1988).

Skeletal mass regressions for *B. musculu*s (r^2^ = 0.910, *n* = 43) and *B. physalus* (r^2^ = 0.856, *n* = 40) were coupled with published soft tissue ratios (Lockyer, 1981) to produce mass models corresponding to the extremes in body conditions (Figures 1, S20, & S21). The ratios for *B. physalus* were updated with additional data (Mogoe et al., 2014). To account for the observed discrepancy in the visceral weight (Omura et al., 1970), adjustments in the visceral components were calculated and refitted into a mass model for *B. m. brevicauda* (Table S3). Figure 1 shows that all *Balaenoptera* species are considerably lighter than balaenids, *E. robustus*, and *M. novaeangliae* at equal lengths. With a 6% correction for body fluids, the regression models predict *B. m. brevicauda* weighing 96.7 tonnes (77.6-115.8 tonnes) at 24.4 m, *B. m. intermedia* weighing 130.7 tonnes (105.0-156.4 tonnes) at 27 m, and 207.8 tonnes (167.0-248.7 tonnes) at 31.1m.

While mass data for *B. musculus* is sparse for individuals exceeding 28 m, observations from oil yields indicate that exceptionally long females did not produce much more than fattened individuals between 24-27.4 m (Walsh, 2010). The record yield was 354 barrels / 56.64 tonnes from a 30.8 m pregnant female while 305 barrels / 48.8 tonnes was extracted from a 27.7 m individual (Tønnessen & Johnsen, 1982). It may be that 27-28 m is the maximum TBL for optimal weight gain and larger whales cannot become especially fat. Oil yields from blubber, meat, and bones compose up to 29% of the total piecemeal mass of a fattened individual (Lockyer, 1981). This predicts an intact body mass of at least 209 tonnes for a whale that produced 56.64 tonnes of oil.

### Relationships between length parameters

The following analyses combined the parameters reported in both the present review and previous work on odontocetes (McClure, 2024). The relationship between *L*_b_, *L*_w,_ and maternal size were previously investigated using interchangeable use of the female *L*^∞^ and *L*_max_ (Huang et al., 2009). This was re-examined using more precise estimates of the female *L*^∞^ and observed values of *L*_b_ (n = 13) and *L*_w_ (n = 12). On average, *L*_b_ equaled 33.6% of *L*^∞^ and was significantly larger in odontocetes (mean = 41.9%, n = 4) than in mysticetes (mean = 29.8%, n = 9, Welch’s two-sample *t*-test: *p* < 0.05). The *L*_w_ averaged 63.5 % of *L*^∞^ and the difference between odontocetes (mean = 73.1%, n = 3) and mysticetes (mean =60.3%, n = 9) was not significant (Welch’s two-sample *t*-test, *p* > 0.05).

Being parameters for adult size, the relationships between *L*_mean_, *L*_sm_, and *L*^∞^ are compared in species and subspecies for which all three were known (Table 10). *E. australis* was omitted due to issues with the compatibility of the *L*^∞^ estimates from photogrammetry data. On average, the *L*_sm_/*L*^∞^, *L*_sm_/*L*_mean_, and *L*_mean_/*L*^∞^ ratios equaled 87.4%, 90.9%, and 96.0%, respectively. None of these values were significantly different between odontocetes and mysticetes (Welch’s two-sample *t*-test, *p* < 0.05). After removing species where < 200 mature individuals were sampled, the species with the highest CV values were *B. mysticetus*, *P. macrocephalus*, and *M. novaeangliae*. As would be expected, the correlation against CV was statistically significant for all three ratios: *L*_sm_/*L*_mean_ (*F*_1,24_=23.1, *p* < 0.001), *L*_sm_/*L*^∞^ (*F*_1,24_= 43.1, *p <* 0.001), and *L*_mean_/*L*^∞^ (*F*_1,24_= 16.4, *p <* 0.001).

**Table 10:**
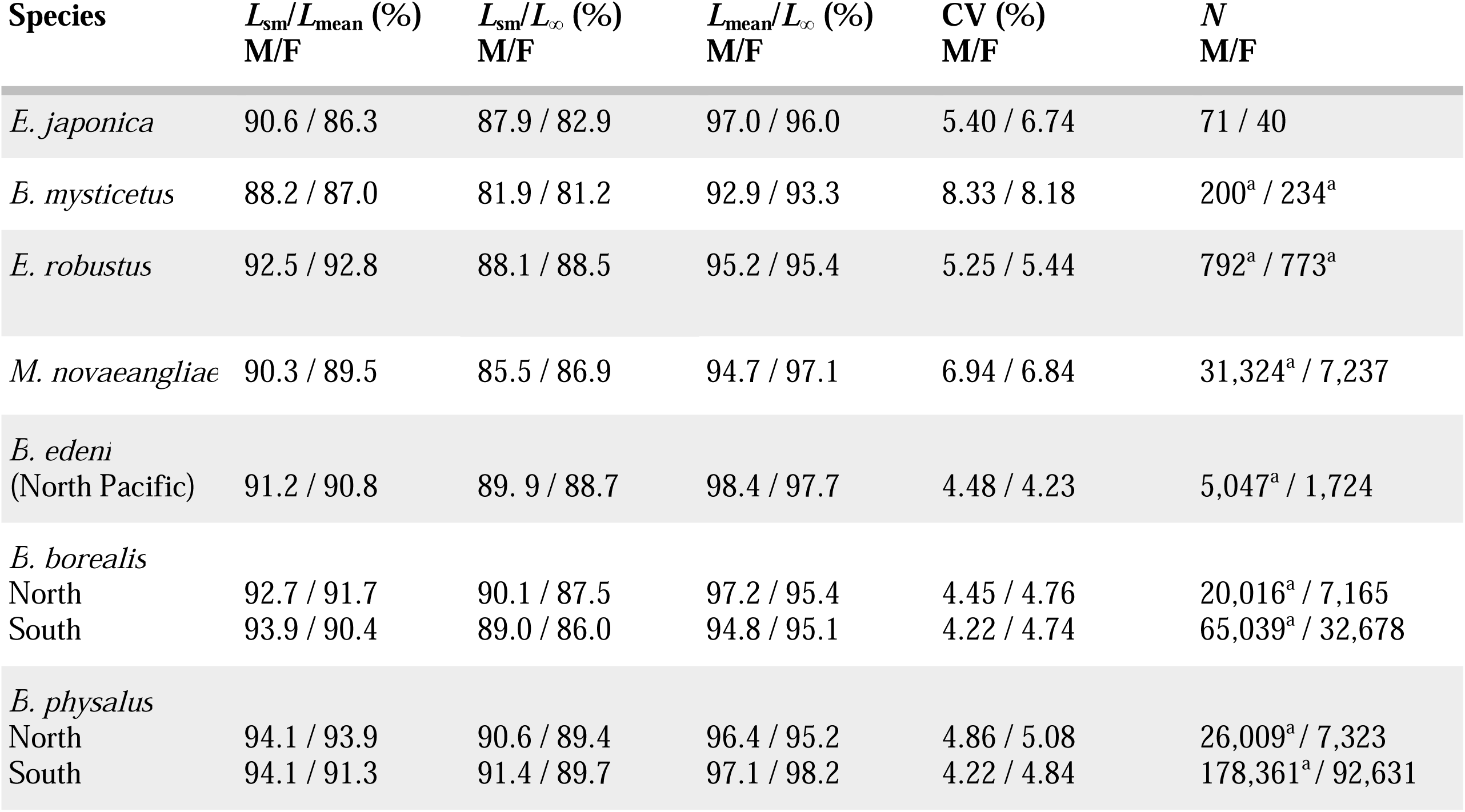
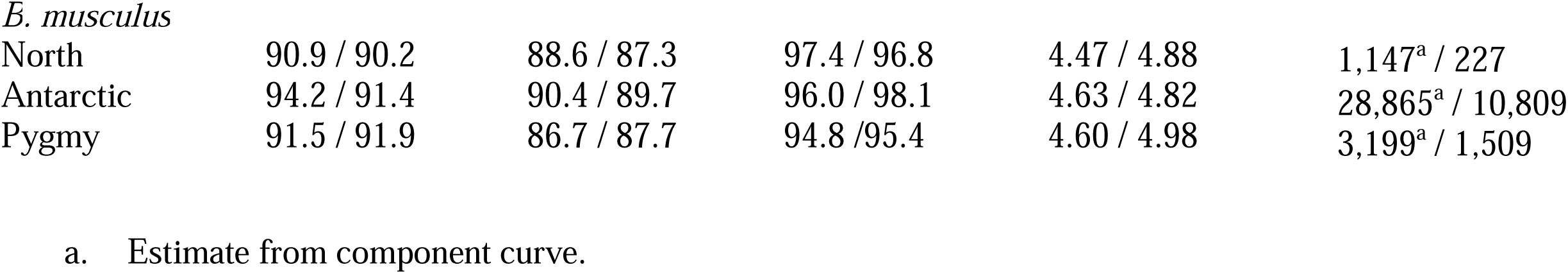
Relationships between *L*_sm_, *L*_mean_, and L_∞_ in large mysticetes.

| Species | $L_{sm}/L_{mean}$ (%)<br>M/F | $L_{sm}/L_{\infty}$ (%)<br>M/F | $L_{mean}/L_{\infty}$ (%)<br>M/F | CV (%)<br>M/F | $N$<br>M/F |
| --- | --- | --- | --- | --- | --- |
| *E. japonica* | 90.6 / 86.3 | 87.9 / 82.9 | 97.0 / 96.0 | 5.40 / 6.74 | 71 / 40 |
| *B. mysticetus* | 88.2 / 87.0 | 81.9 / 81.2 | 92.9 / 93.3 | 8.33 / 8.18 | 200<sup>a</sup> / 234<sup>a</sup> |
| *E. robustus* | 92.5 / 92.8 | 88.1 / 88.5 | 95.2 / 95.4 | 5.25 / 5.44 | 792<sup>a</sup> / 773<sup>a</sup> |
| *M. novaeangliae* | 90.3 / 89.5 | 85.5 / 86.9 | 94.7 / 97.1 | 6.94 / 6.84 | 31,324<sup>a</sup> / 7,237 |
| *B. edeni*<br>(North Pacific) | 91.2 / 90.8 | 89.9 / 88.7 | 98.4 / 97.7 | 4.48 / 4.23 | 5,047<sup>a</sup> / 1,724 |
| *B. borealis*<br>North | 92.7 / 91.7 | 90.1 / 87.5 | 97.2 / 95.4 | 4.45 / 4.76 | 20,016<sup>a</sup> / 7,165 |
| South | 93.9 / 90.4 | 89.0 / 86.0 | 94.8 / 95.1 | 4.22 / 4.74 | 65,039<sup>a</sup> / 32,678 |
| *B. physalus*<br>North | 94.1 / 93.9 | 90.6 / 89.4 | 96.4 / 95.2 | 4.86 / 5.08 | 26,009<sup>a</sup> / 7,323 |
| South | 94.1 / 91.3 | 91.4 / 89.7 | 97.1 / 98.2 | 4.22 / 4.84 | 178,361<sup>a</sup> / 92,631 |
| *B. musculus* |  |  |  |  |  |
| North | 90.9 / 90.2 | 88.6 / 87.3 | 97.4 / 96.8 | 4.47 / 4.88 | 1,147<sup>a</sup> / 227 |
| Antarctic | 94.2 / 91.4 | 90.4 / 89.7 | 96.0 / 98.1 | 4.63 / 4.82 | 28,865<sup>a</sup> / 10,809 |
| Pygmy | 91.5 / 91.9 | 86.7 / 87.7 | 94.8 / 95.4 | 4.60 / 4.98 | 3,199<sup>a</sup> / 1,509 |

a. Estimate from component curve.

## 4 Discussion

The discrepancy between commercial data and photogrammetry measurements for *E. australis* mirrored previous observations for both this species (Tormosov et al., 1998) and *B. mysticetus* (George et al., 2004), where hauled whales appeared to be systematically longer. This phenomenon is observed to neither the same degree nor consistency in more streamlined species that overlap in body mass (Bierlich et al., 2023; Miller et al., 2013; Ortega-Ortiz et al., 2022; Russell et al., 2023, 2024). Rather than the hauling process uniformly stretching the tissue in the carcasses of large whales, the discrepancy appears to reveal that the more rotund balaenids experiencing a greater post-mortem deviation from their natural posture while lying upon a flat surface.

Across the Southern Hemisphere dataset for *B. physalus,* several expeditions reported mature females as small as 14.3-16 m. This is notable as the smallest mature females reported in biological studies typically measured 18.5-18.6 m (Kakuwa et al., 1953; Mackintosh, 1942; Ohno & Fujino, 1952). While unusually small mature females would align with the observations made for a proposed pygmy subspecies (Clarke, 2004), clerical errors or species misidentification must also be considered.

For *B. musculus*, the SE Pacific catch was most similar in mean size to the northern subspecies, in agreement with the results of genetic clustering with the ENP stock (Attard et al., 2024). The size difference from the pooled North Pacific catch data aligns with previous findings in separating the ENP and WNP populations (Monnahan et al., 2014). The South Pacific catch for *B. borealis* mirrors this pattern without any obvious signs of species misidentification with *B. edeni*. While biological studies do not seem to be available, the acoustic data from *B. borealis* near Chile were unique from all other regions (Español-Jiménez et al., 2019). This would support that the SE Pacific stock for *B. borealis* may be a discrete population from the rest known in the Southern Hemisphere.

Interspecific differences in body mass were notably grouped by feeding strategy: lunge-feeding *Balaenoptera* were the lightest and the skim-feeding balaenids were the heaviest by a substantial margin, as previously acknowledged (George, 2009; Lockyer, 1976). Intermediate of the two groups were *E. robustus* and *M. novaeangliae*, generalists that employ a combination of different feeding mechanisms (Croll et al., 2018; Nerini, 1984). Given the high cost of lunge-feeding (Potvin et al., 2012), it is likely that the different mass relationships and overall morphology of these species reflect the combination of feeding energetics and prey availability. Due to limited sampling, weight variation remains understudied for certain stocks, such as those in the North Atlantic for most species. This may be resolved as the methodology of photogrammetric body mass estimates continue to advance. These techniques also offer the advantage of performing longitudinal comparisons for the same individuals.

It is notable that photogrammetric volume and mass models for *B. musculus*, *M. novaeangliae*, *E. robustus, P. macrocephalus* conform to average body tissue densities near to that of seawater (Bernier-Graveline et al., 2025; Christiansen et al., 2021, 2025; Glarou et al., 2022; Napoli et al., 2024; Russell et al., 2023, 2024; van Aswegen, 2025). By contrast photogrammetric models for balaenids, when calibrated from mass data from hauled whales, predicted notably lower body density estimates (Christiansen et al., 2019, 2024). As the true body mass remains constant as the posture changes, the volume of hauled whales might be overestimated by models designed for photographed whales in their natural posture. This would then downsize the average body density of landed whales with overestimated volumes (Christiansen et al., 2019, 2024). This interpretation may be further validated through more direct body density estimates for balaenids, such as direct sampling of tissue (van Aswegen, 2025) or possibly hydrodynamic gliding models (Aoki et al., 2021).

Analyses of early growth parameters re-affirmed the convergence in relative size at weaning while birth size in odontocetes was found to be relatively greater than that of mysticetes. This is consistent with odontocete gestation generally lasting about 50% longer (Best et al., 1984; Kasuya, 1977; Lockyer, 1984; Olesiuk et al., 2005). As capital breeders, mysticetes accumulate energy stores that foster considerably faster growth during lactation than the continuous reserves of odontocetes (Irvine et al., 2017; Oftedal, 1997). Odontocetes producing proportionally larger neonates through longer gestations may serve to compensate for the slower postnatal growth towards weaning size. This could be an important factor to explore in future research on the adaptational divergence in cetacean reproduction.

Without direct data from biological examinations or photogrammetric surveys, size parameters for certain cetacean species are estimated using other species as analogs (Huang et al., 2009; Pastene et al., 2020). In general, the *L*_mean_ for large cetaceans was ∼4% lower than the *L*^∞^, and the CVs were about 4-5% in most species. *P. macrocephalus*, *M. novaeangliae*, and *B. mysticetus* having the highest variability in adult size (6-8%) is consistent with previous research indicating more prolonged growth after sexual maturity and very low frequencies of physically mature individuals (Chittleborough, 1955b, 1955a; Clarke et al., 1994; Lubetkin et al., 2012). Being able to use the variation in adult size to infer the interval of adult growth and vice-versa can have important applications in modeling the growth and size distributions of under-studied baleen whale stocks or fossil species (Lambert et al., 2010).

## 5 Conclusions

Previous reviews have attempted to provide accurate descriptions for the size of some large mysticete species (McClain et al., 2015; Wood, 1982), but were limited in either their accuracy or species coverage. Prioritizing primary literature has revealed several instances where the republished data was subject to typographical errors or were missing significant details addressed by the original authors (Klumov, 1962; Scheffer, 1974; Voronin, 1948).

This review on the size of large cetaceans has compiled a large breadth of morphometric parameters for postnatal growth, adult variation, and body mass relationships. Analyzing the information from both the literature and the IWC catch database facilitated descriptions of the variance and maximum credible sizes for the current species. This review will serve as an excellent reference for future research and provide another cornerstone in advancing morphometric analysis that emphasizes the importance of variability within species.

## Data availability

Supplementary figures, code, and morphometric data acquired from literature and museum records can be acquired upon contact with the corresponding author. All whaling data is available from the International Whaling Commission upon request to this address

IWC contact page: https://iwc.int/contact

## Acknowledgements

The author expresses appreciation towards Dr. Sally Mizroch, Dr. Trevor Branch, Dr. Fredrik Christiansen, Dr. Yulia Ivashchenko, Dr. Phil Clapham, Dr. James Mead, Seán O’Callaghan, and Samuel Dunford for both their correspondence and assistance in acquiring important sources for this paper.

