## Supplementary File 1 for "A summary of intraspecific size variation for large mysticetes"

### **A summary of intraspecific body size variation of large mysticetes. Supplementary File 1**

Author: Joseph McClure*

*Correspondence

564 East McIver Road, Florence, South Carolina 29506

#### Figure S1. Female (A) and male (B&C) length distributions and normal probability plot for *E. australis* from Soviet catch data with fitted curves for mature whales (approx. 81% of males were mature). Mature female (D) and male (E) length distributions for *E. japonica*. See Supplementary file 2 for *E. japonica* data.

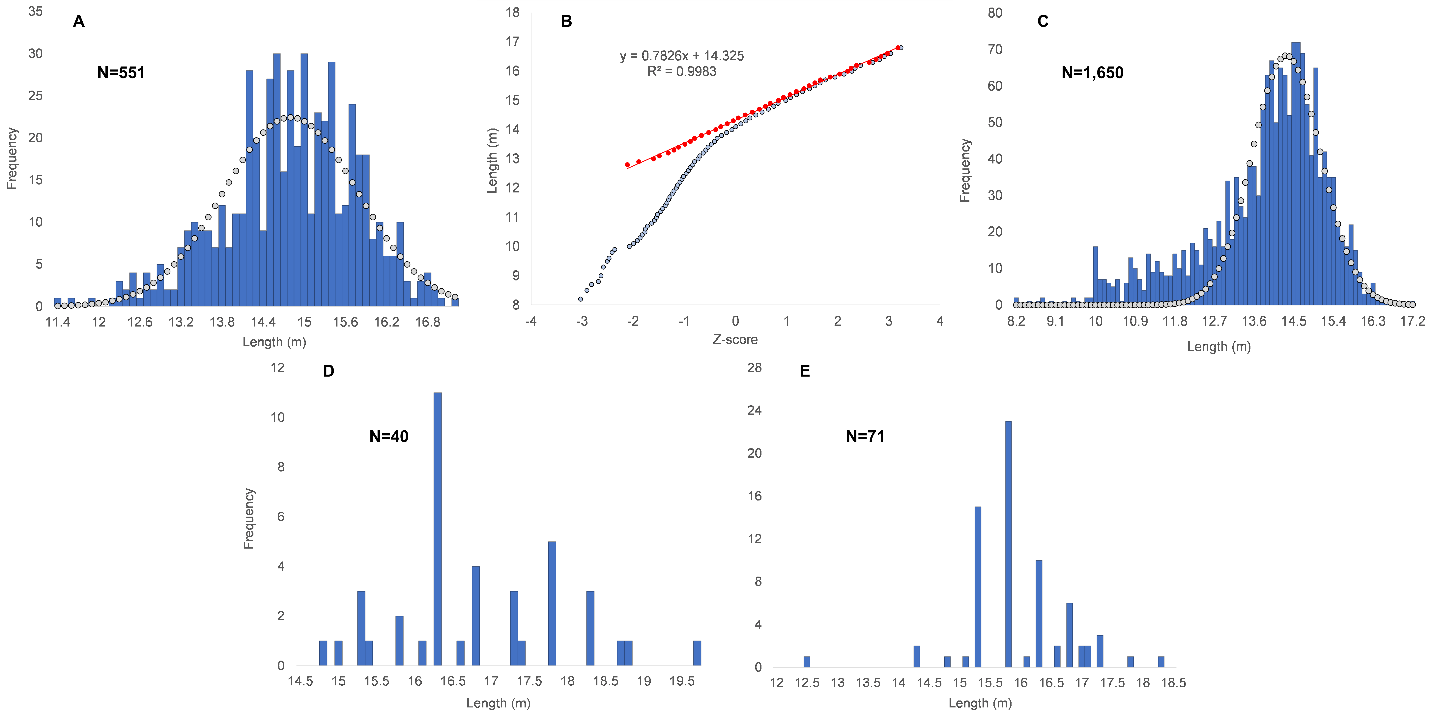

#### Figure S2. Female (A&B) and male (C&D) normal probability plots and length distributions for *B. mysticetus* with fitted curves for mature whales. Approximately 30% of males and 32% of females were mature.

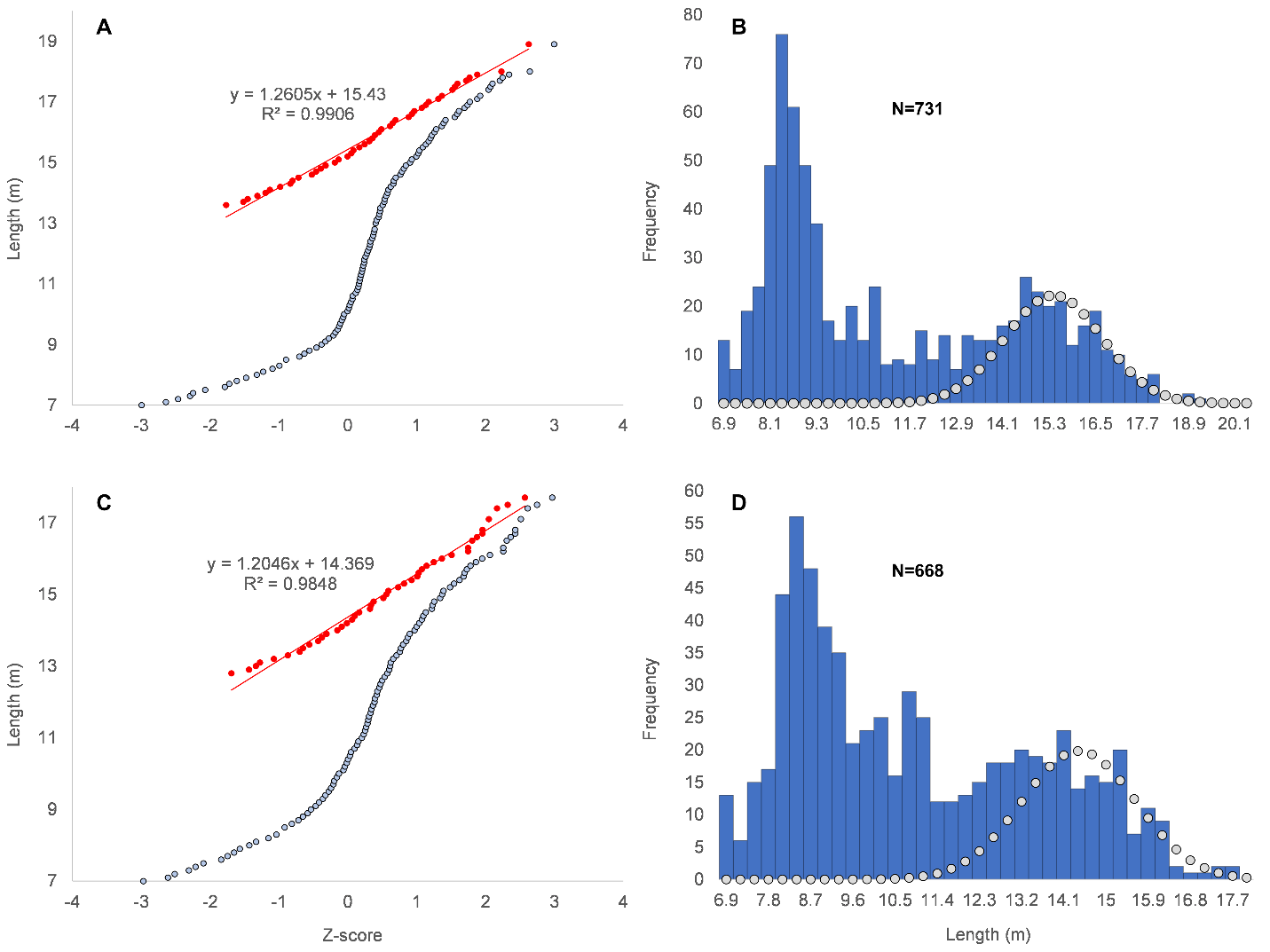

#### Figure S3. (A) Linear regression for weight data on Balaenidae (y=3.129x-1.952, n=39, r^2^= 0.986). Two outliers from necropsy data (Fortune et al., 2021) were omitted from regression. (B) Comparison of Rice-Wolman models against mass data for Balaenidae. Model RW1 uses integer parameters while RW2 uses non-integer parameters. Supplementary 2 for appendix and File 3 for sources.

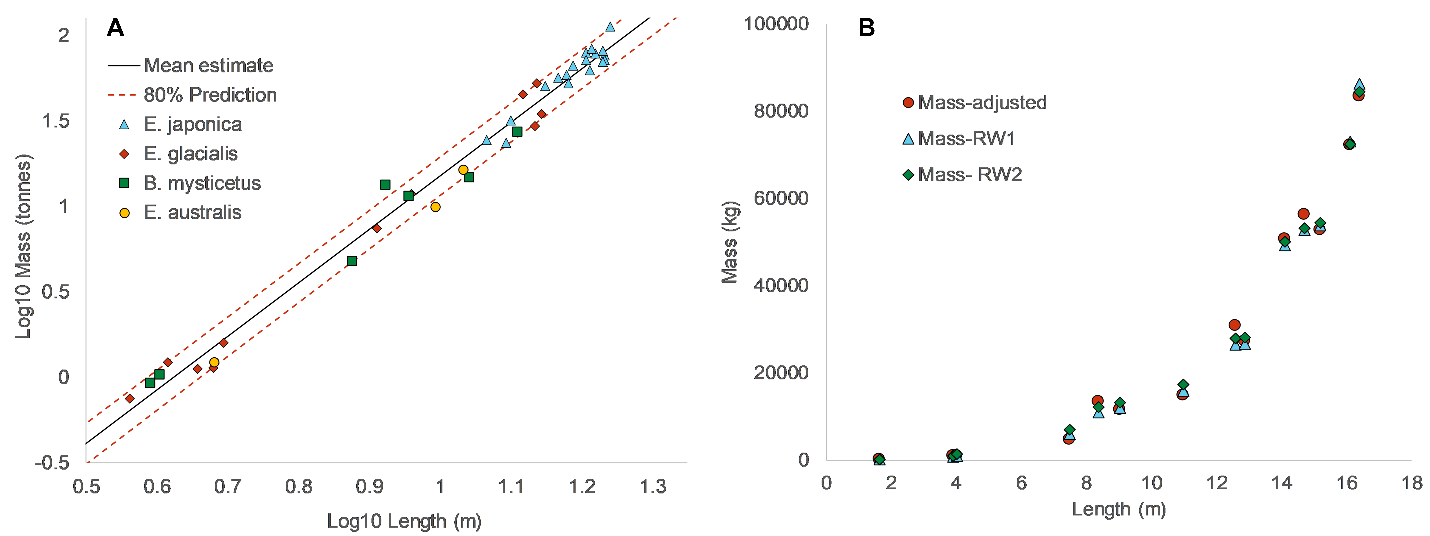

#### Table S1. Weight model parameters for Balaenids.

| Formula | Species | a | b/b1 | b2 | n | Units  (length / mass) | Reference |
| --- | --- | --- | --- | --- | --- | --- | --- |
| Eq. 2 | *E. japonica* | 0.0124 | 3.06 | — | 21 | m / tonnes | Lockyer, 1976 |
| Eq. 2 | *E. australis* | 0.0131^a^ | 3.016 | — | 5,372 | m / tonnes | Christiansen et al., 2022 |
| Eq. 2 | *E. glaclias /*  *E. japonica* | 0.0114^b^ | 3.06 | — | 29 | m / tonnes | Fortune et al., 2021 |
| Eq. 2 | Balaenid sp*.* | 0.0112^c^ | 3.129 | — | 39 | m / tonnes | Current study |
| Eq. 3 | *B. mysticetus* | 38.53^c^ | 1 | 2 | 8 | m / kg | George, 2009 |
| Eq. 3 | Balaenid sp. | 38.7^c,d^ | 1 | 2 | 14 | m / kg | Current study |
| Eq. 3 | Balaenid sp. | 58.9^c,d^ | 0.95 | 1.87 | 14 | m / kg | Current study |

a. Coefficient when average body density is set to 805 kg/ m^3^.
b. Value converted for meters and tonnes.
c. Includes fluid adjustments and Case Nos. 32 & 45 from Fortune et al., 2021 were excluded.
d. Pregnant individuals excluded.

#### Figure S4. Female (A&B) and male (C&D) normal probability plots and length distributions for *E. robustus*. Approximately 43% of males and 33% of females were mature.

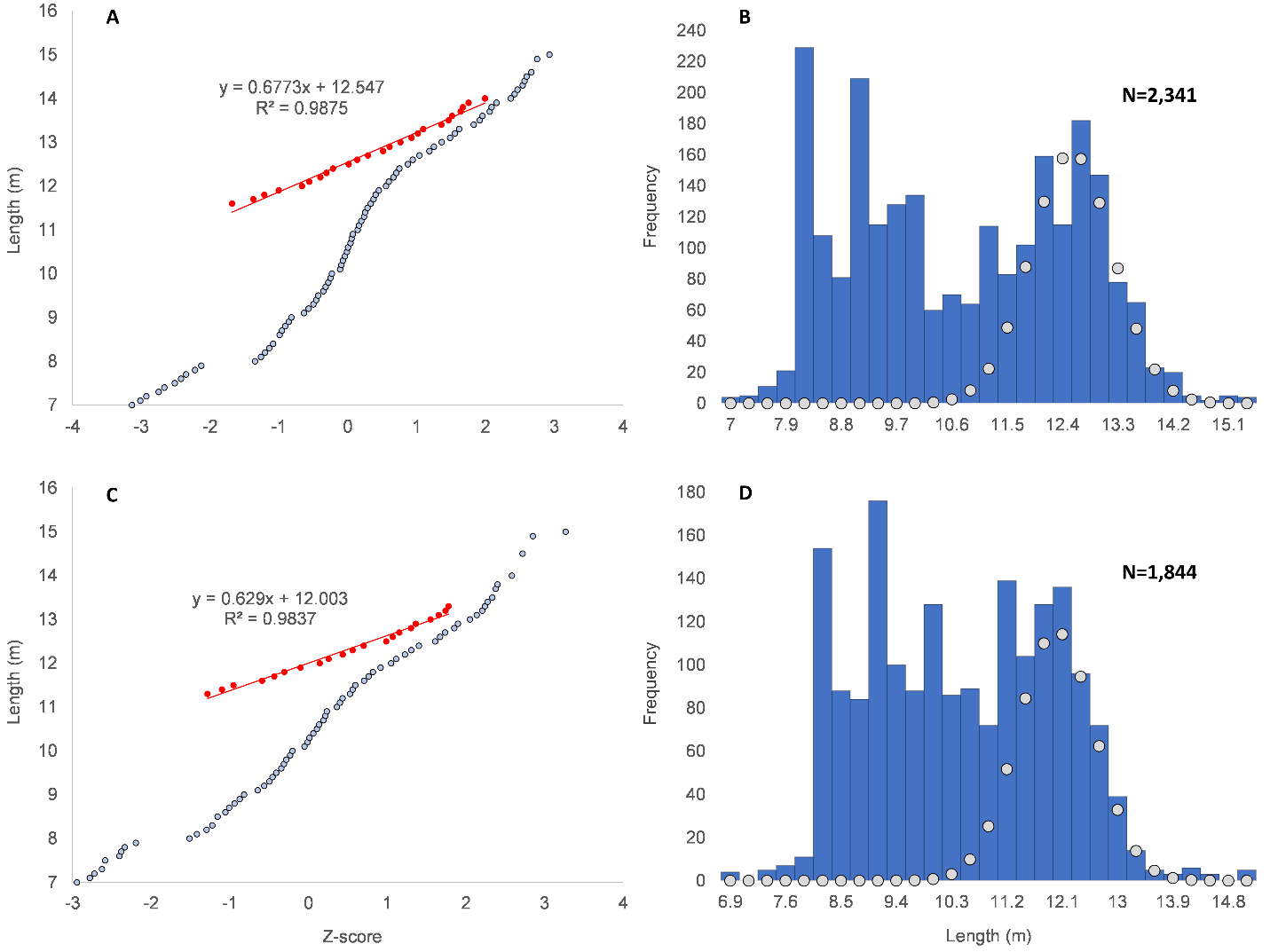

#### Table S2. Weight model parameters for *E. robustus*

| Formula | a | b/b1 | b2 | n | Units (length / mass) | Reference |
| --- | --- | --- | --- | --- | --- | --- |
| Eq. 3 | 20.9^a^ | 2 | 1 | 2 | m / kg | J. Sumich, 1986 |
| Eq. 3 | 28.5^b^ | 1.73 | 1.17 | 12 | m / kg | J. Sumich et al., 2013 |
| Eq. 2 | 9.76^b^ | 3.05 | — | 14 | m / kg | J. Sumich et al., 2013 |
| Eq. 2 | 0.0051 | 3.28 | — | 8 | m / tonnes | Lockyer, 1976 |
| Eq. 2 | 0.01085^c^ | 2.9509 | — | 15 | m / tonnes | Agbayani et al., 2020 |
| Eq. 2 | 0.04529 | 2.570 | — | 288 | m / tonnes | Christiansen et al., 2024 |

a. Parameter for pregnant whales
b. Fitted to measurements taken from live-weighings of captive specimen, JJ.
c. Value converted to meters and tonnes

#### Figure S5. Female (A) and male (B&C) length distributions and normal probability plot from the global catch of *M. novaeangliae* with fitted curves for mature whales. Approximately 78% of males were mature.

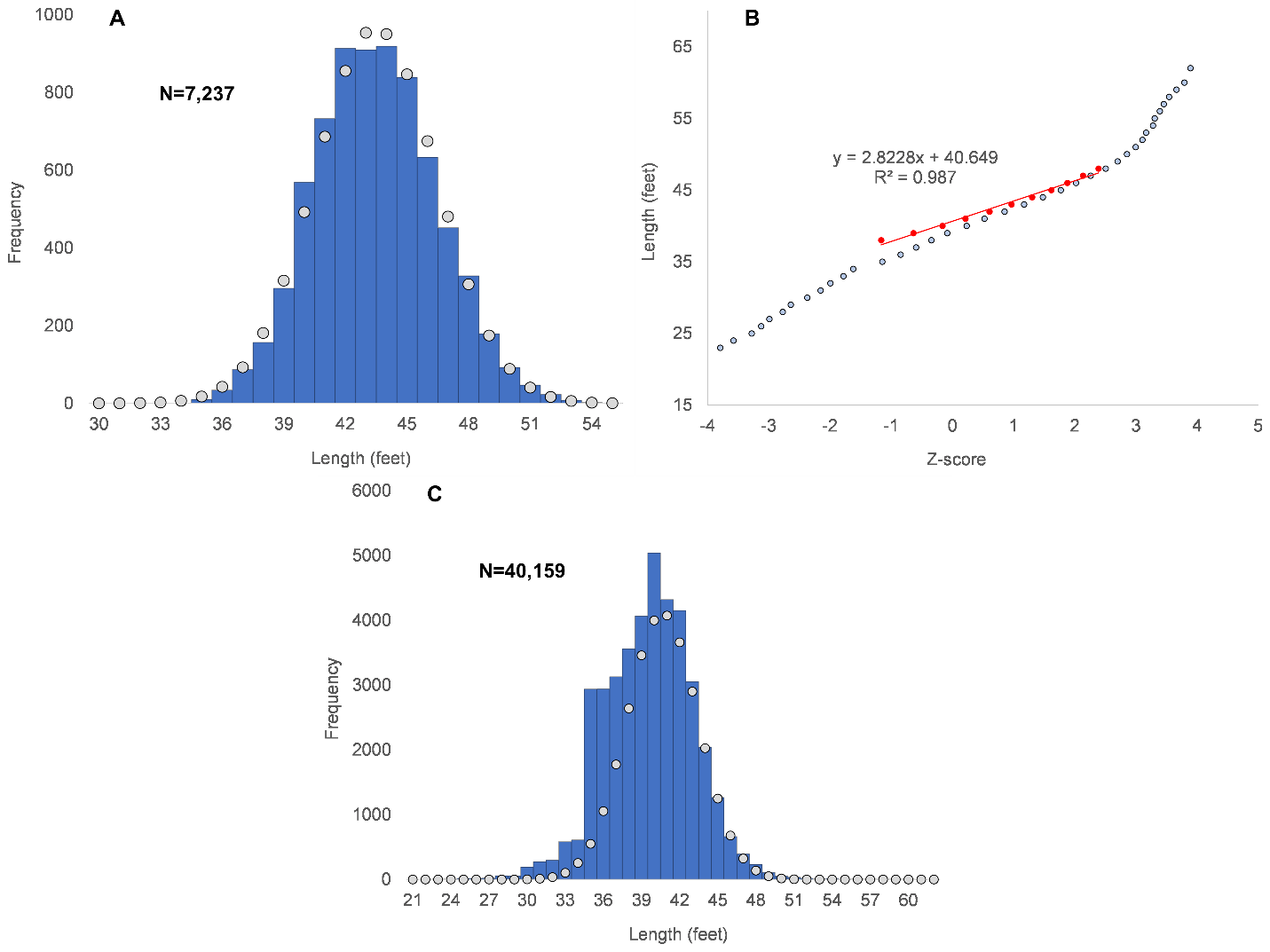

#### Figure S6. Female *B. edeni* length distributions with fitted curves for mature whales from the North Pacific (A), Southern Hemisphere Pelagic catch (B), South Atlantic (C), Southern Hemisphere Pelagic-Soviet (D).

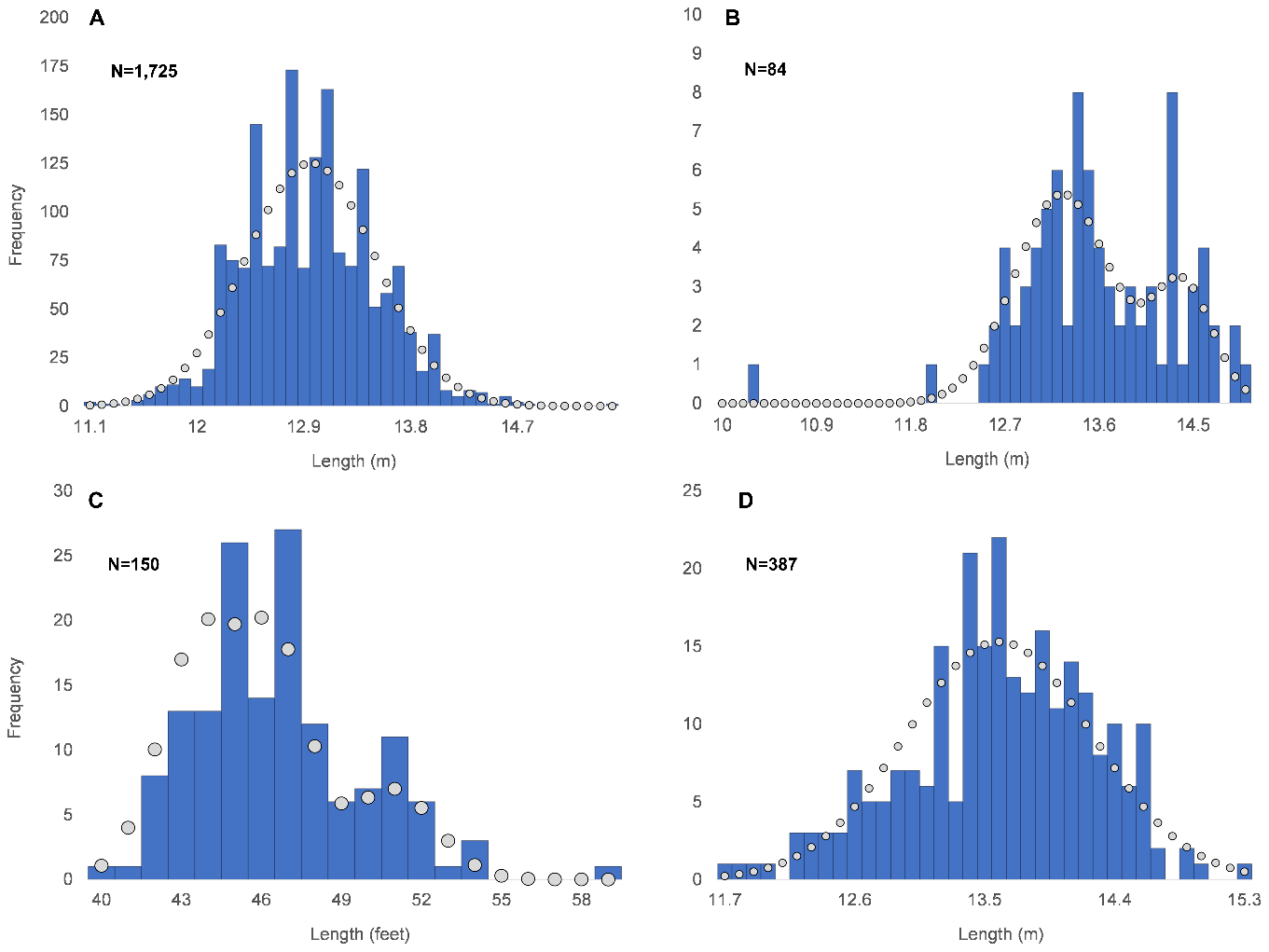

#### Figure S7. Female *B. edeni* normal probability plots for mature whales from the South Atlantic (A) and Southern Hemisphere Pelagic catches (D). The South Atlantic catch contains a probable mixture of *B. borealis* (red line, 19%), offshore morph (orange line, 30%) and inshore morph whales (blue line, 51%). Pelagic data shows a mixture of a larger (red line, 26%) and smaller morphs (blue line, 74%).

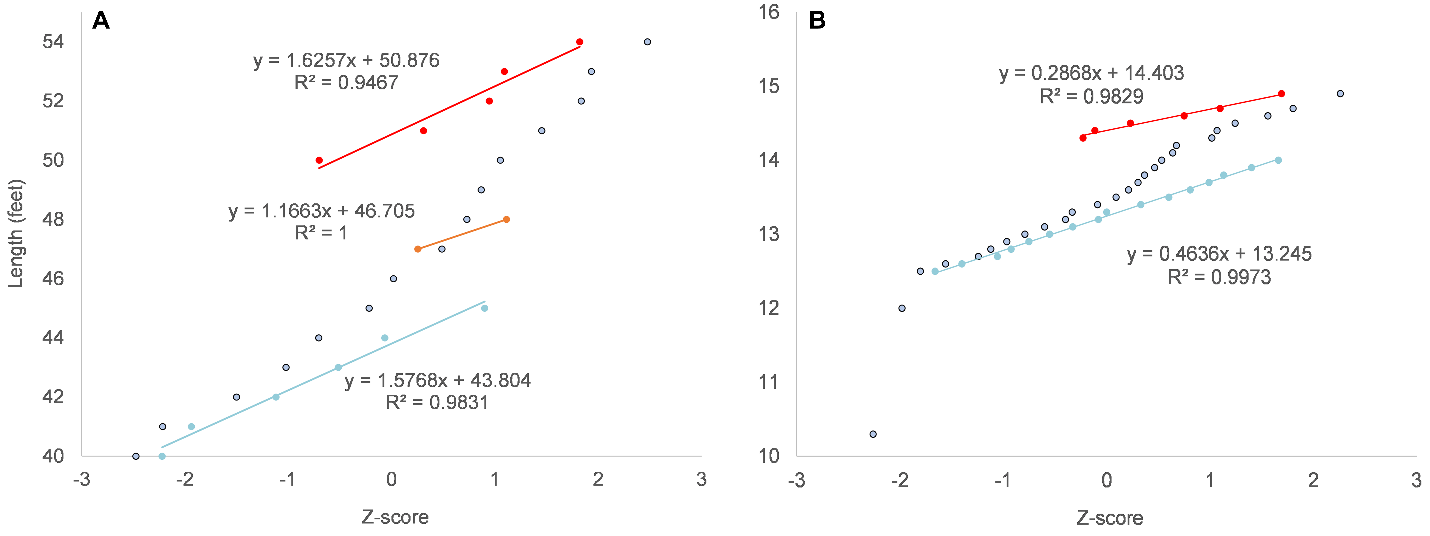

#### Figure S8. Male *B. edeni* length distributions with fitted curves for mature whales from the North Pacific (A), Southern Hemisphere Pelagic (B), South Atlantic (C), Southern Hemisphere Pelagic-Soviet (D) catches. The approximate number of mature individuals are listed in Table 7.

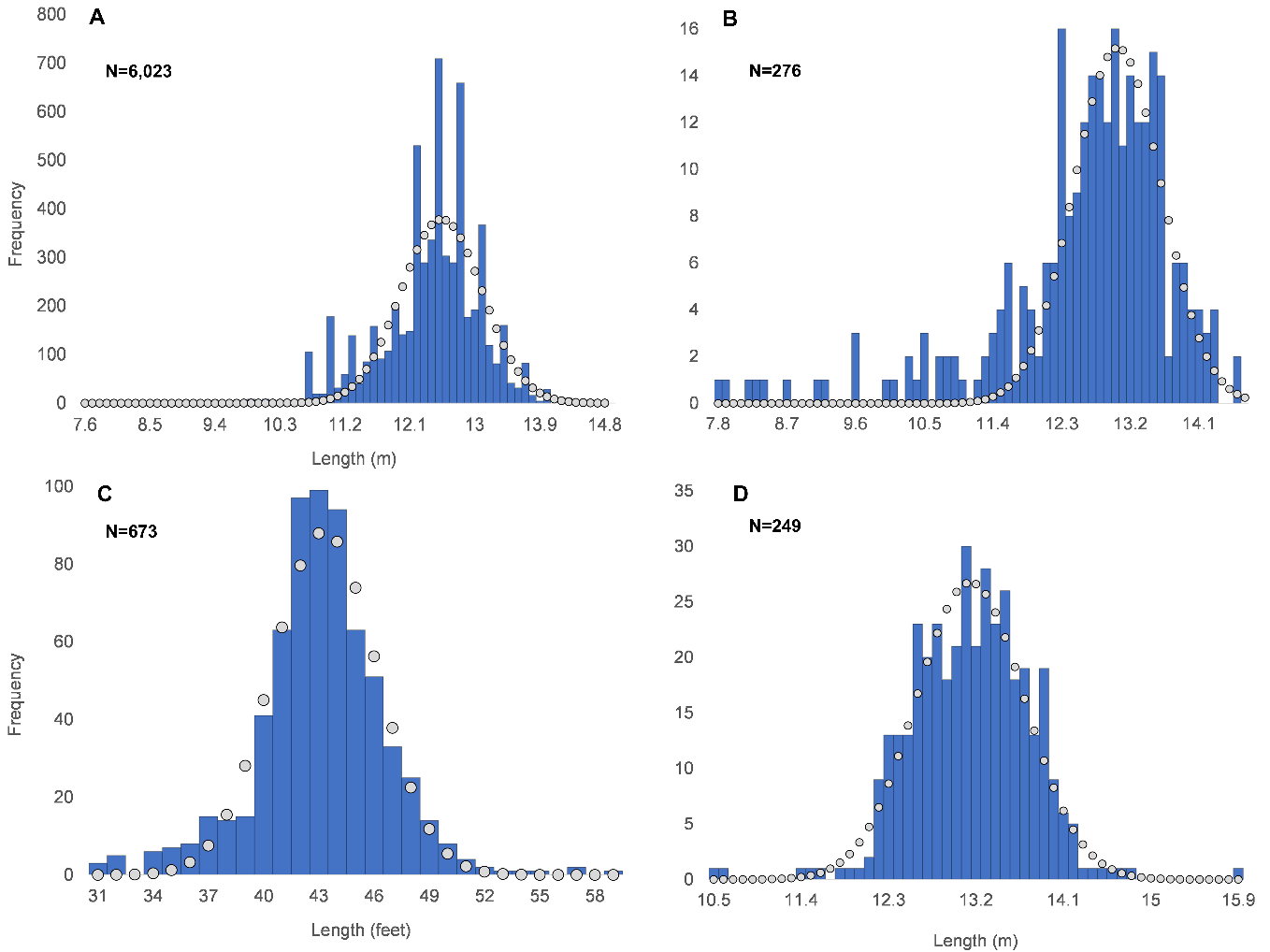

#### Figure S9. Male *B. edeni* normal probability plots for the North Pacific (A), South Atlantic (B), Southern Hemisphere Pelagic (C), Soviet Southern Hemisphere Pelagic catches (D).

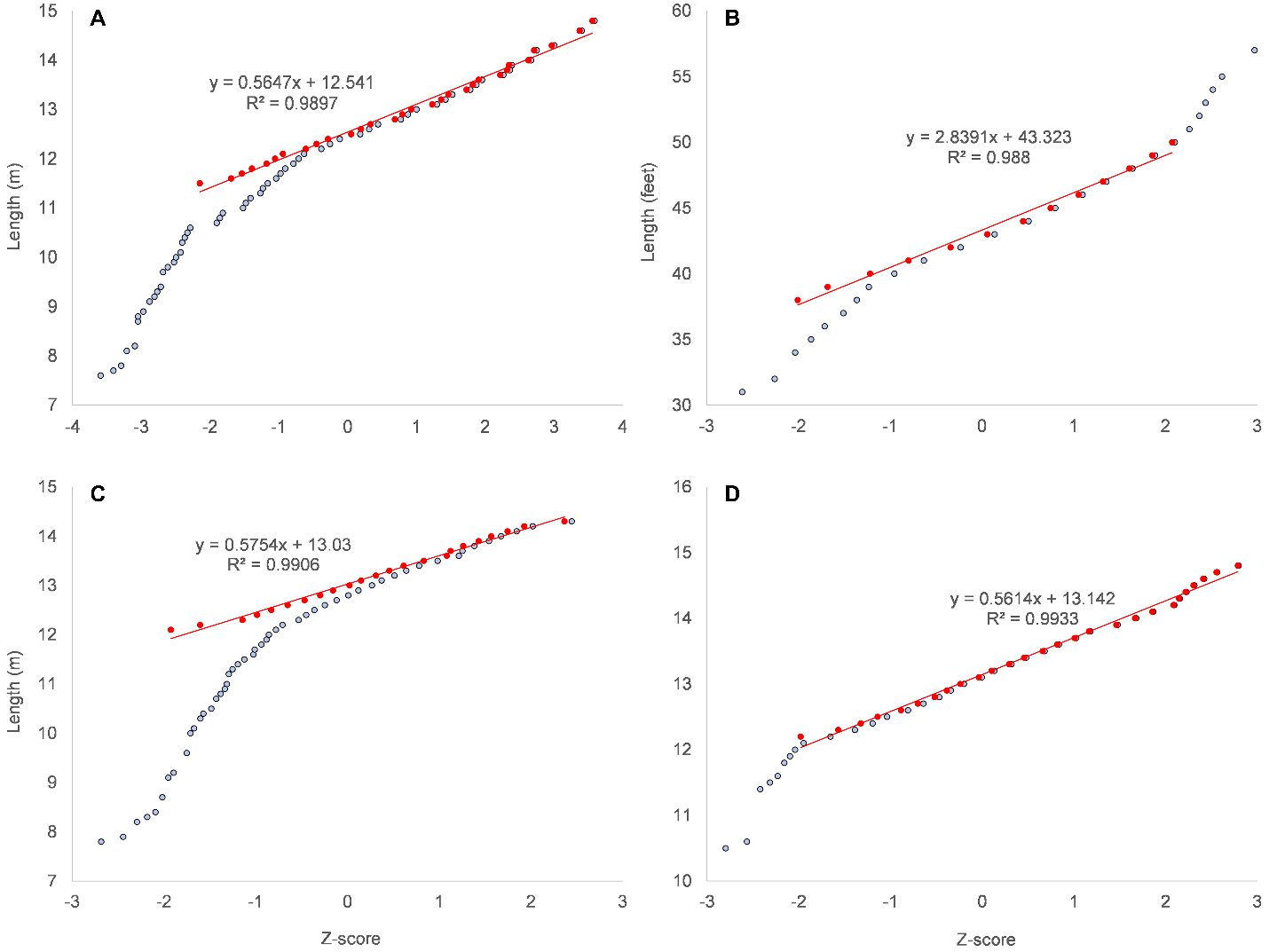

#### Figure S10. Female *B. borealis* length distributions with fitted curves for mature whales in the Northern Hemisphere (A) and Southern Hemisphere (B) and South Pacific (C).

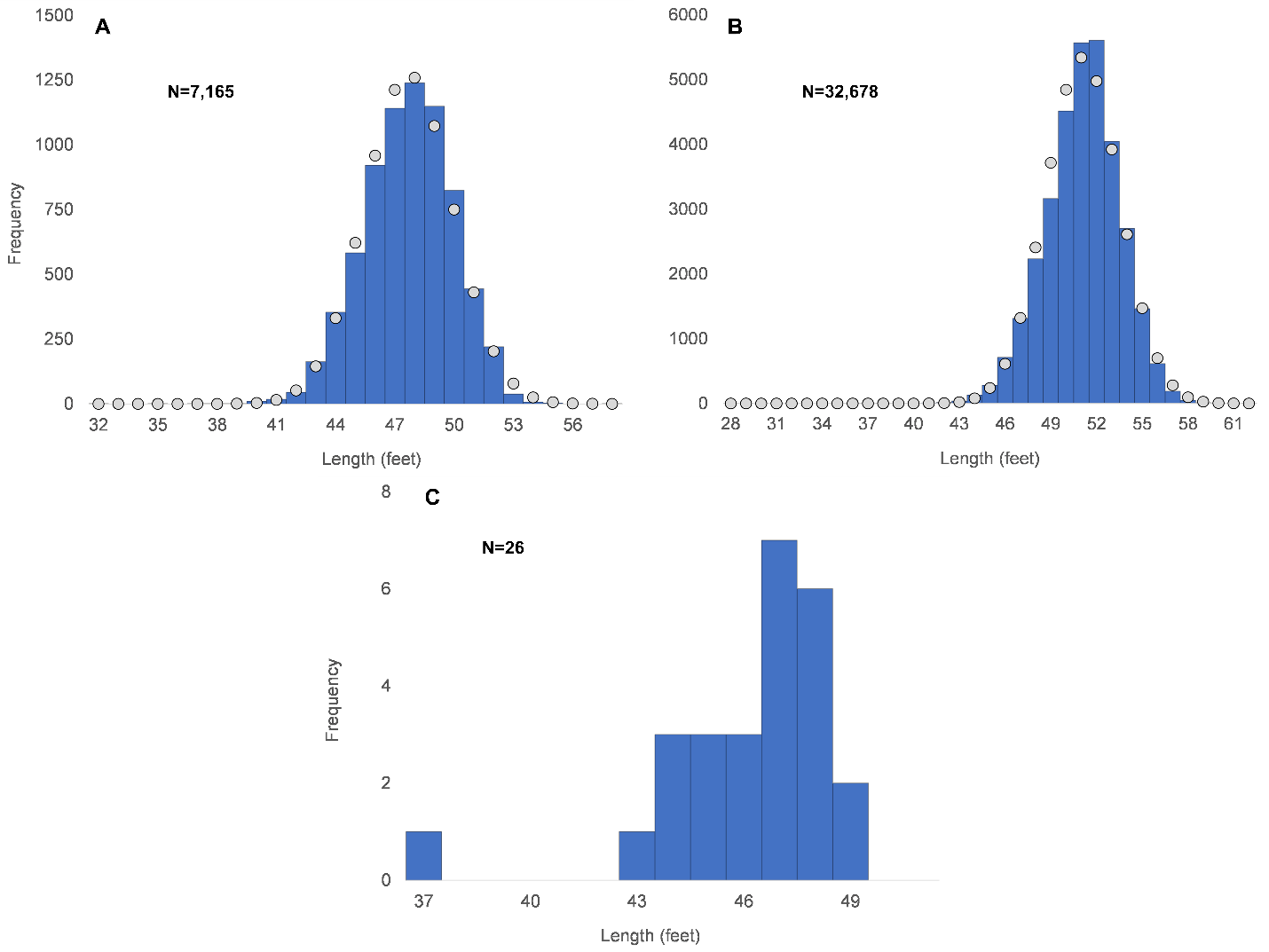

#### Figure S11. Male *B. borealis* normal probability plots and length distributions with fitted curves for mature whales in the Northern (A&B) and Southern (C&D) Hemispheres. Approximately 73% of northern males and 77% of southern males were mature.

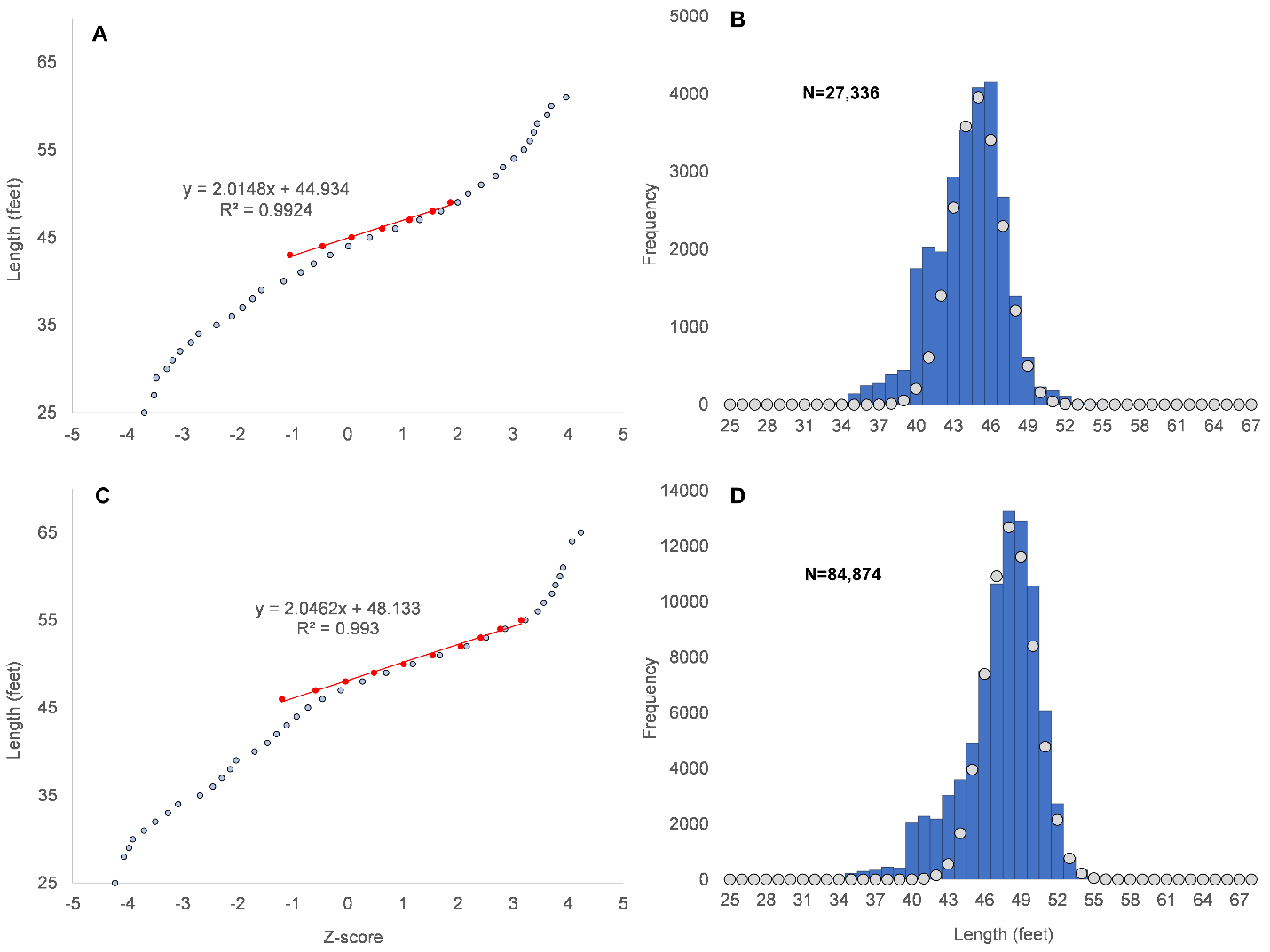

#### Figure S12. Female *B. physalus* normal probability plots and length distributions with fitted curves for mature whales in the Northern (A) and Southern (B) Hemispheres.

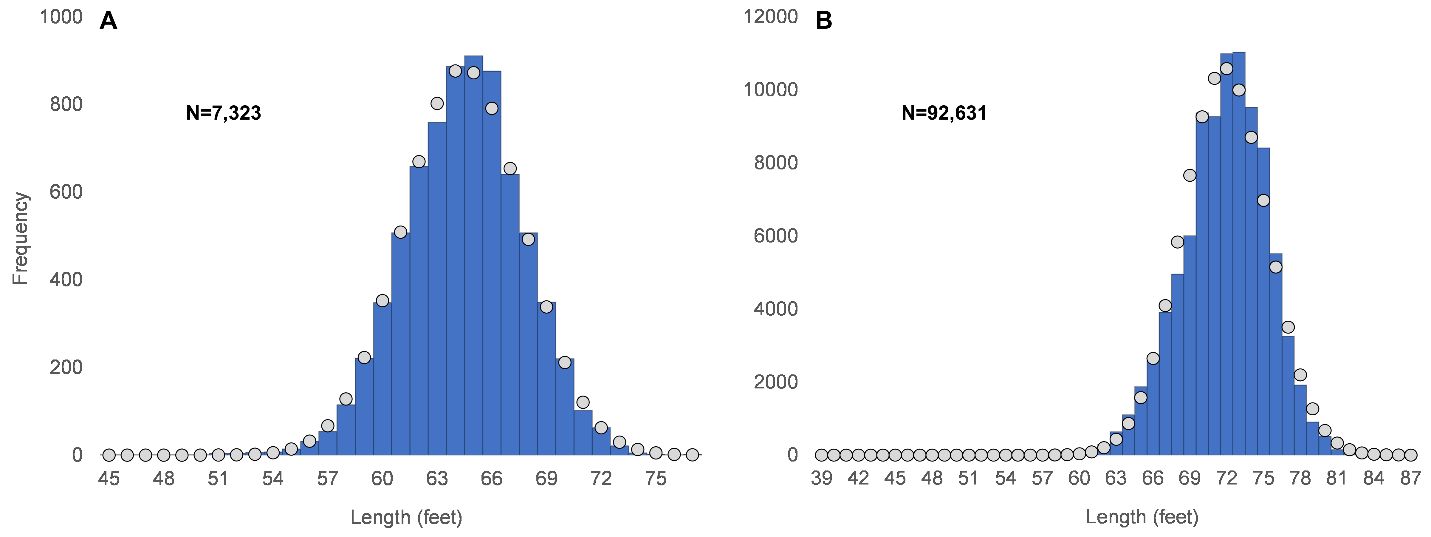

#### Figure S13. Female *B. physalus* length distribution for mature whales from the South Atlantic, South Pacific, and Indian Ocean catches.

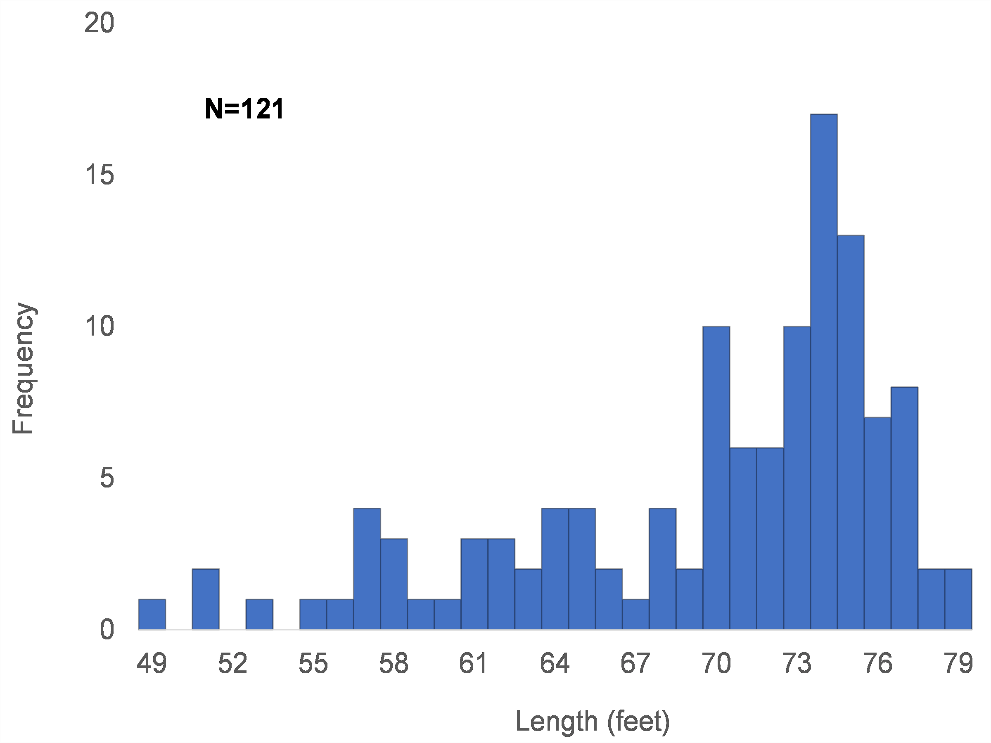

#### Figure S14. Male *B. physalus* normal probability plots and length distributions with fitted curves for mature whales in the Northern (A&B) and Southern (C&D) Hemispheres. Approximately 75% of northern males and 68% of southern males were mature.

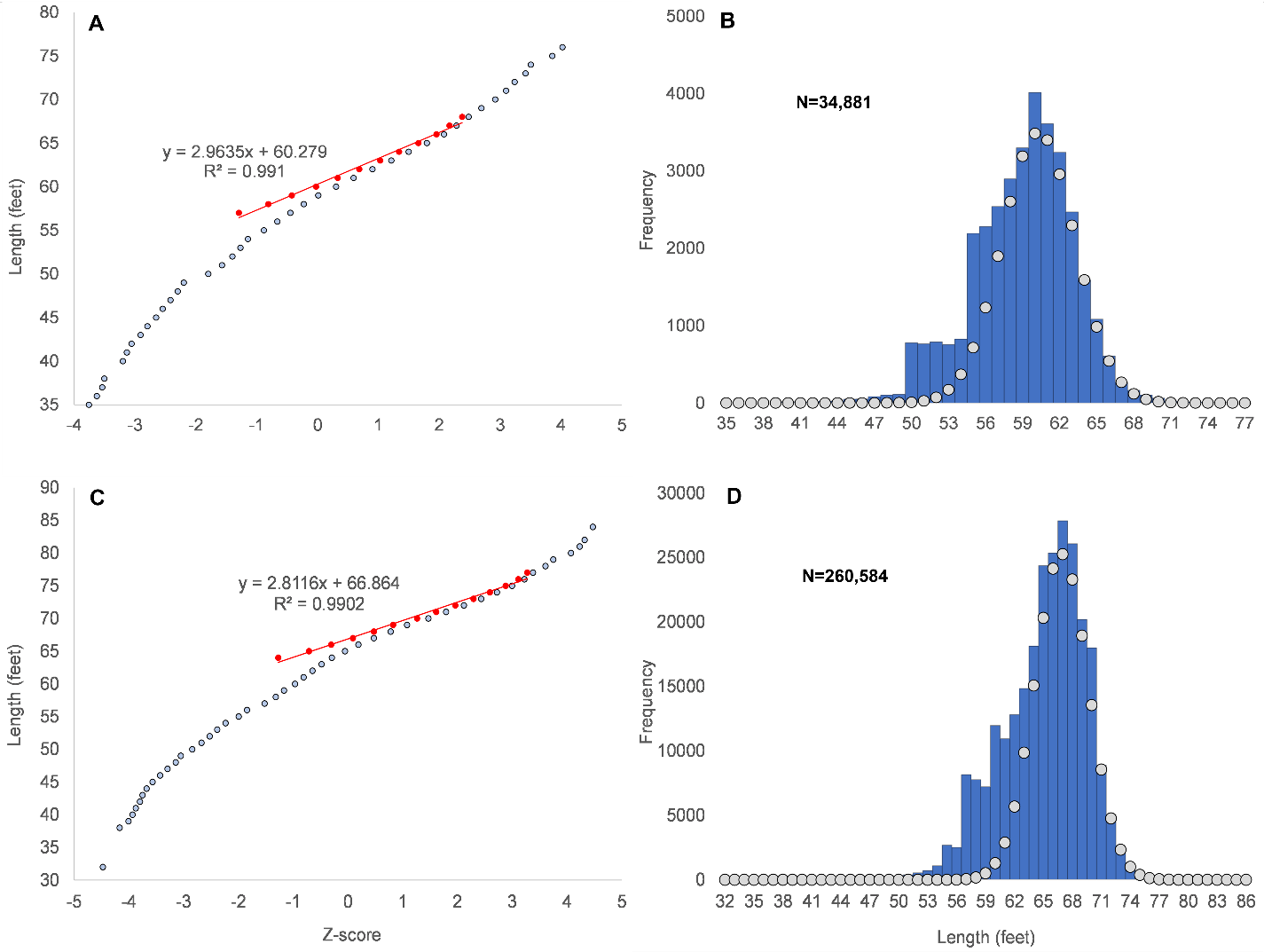

#### Figure S15. Female *B. musculus* length distributions with fitted curves for mature whales from the North Pacific (A), North Atlantic (B), Antarctic (C), Chilean (D), and Pygmy catches (E).

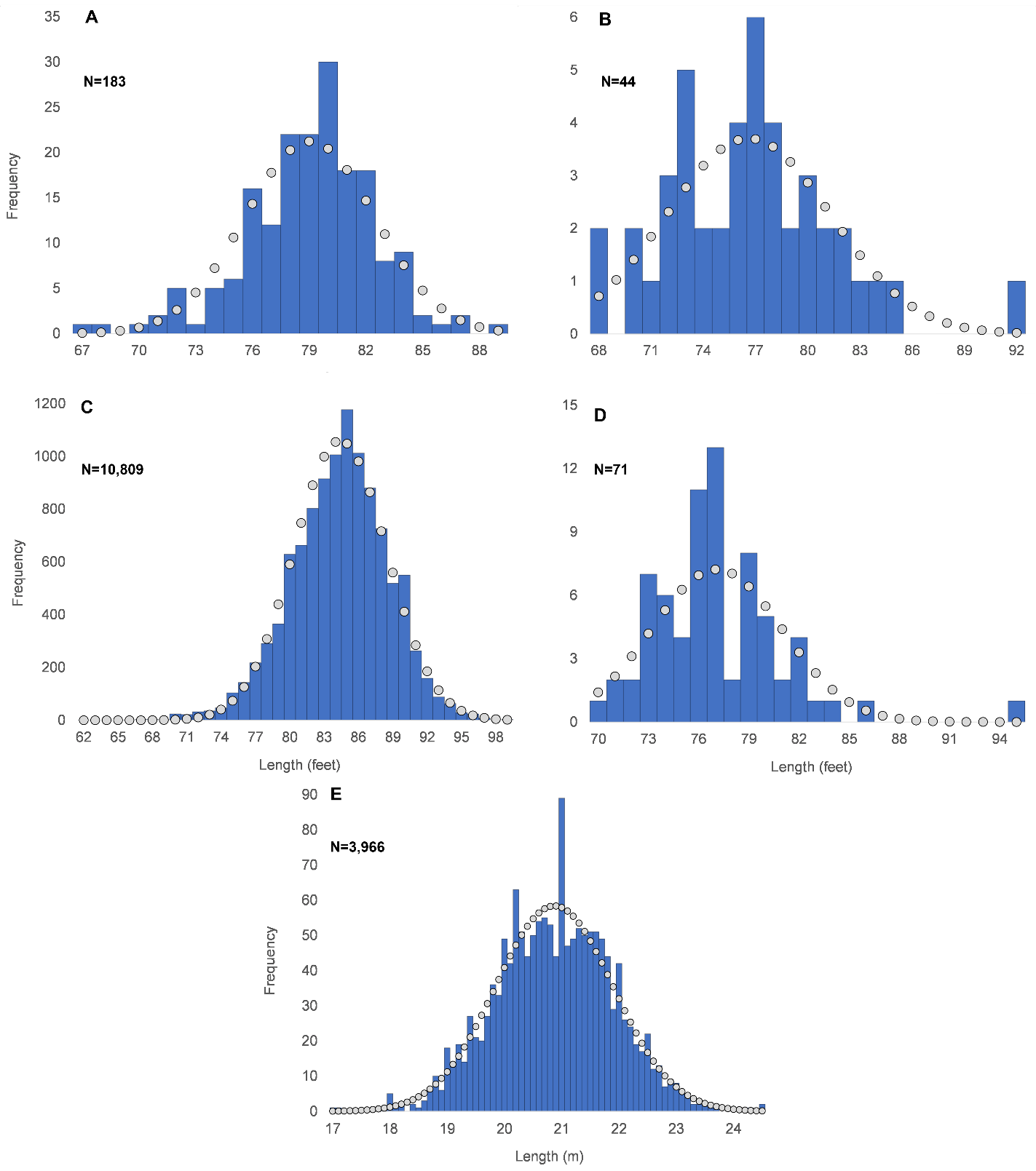

#### Figure S16. Male *B. musculus* length distributions with fitted curves for mature whales from the North Pacific (A), North Atlantic (B), Antarctic (C), Chilean (D), and Pygmy (E) catches. Approximate number of mature individuals for males are listed in Table 10.

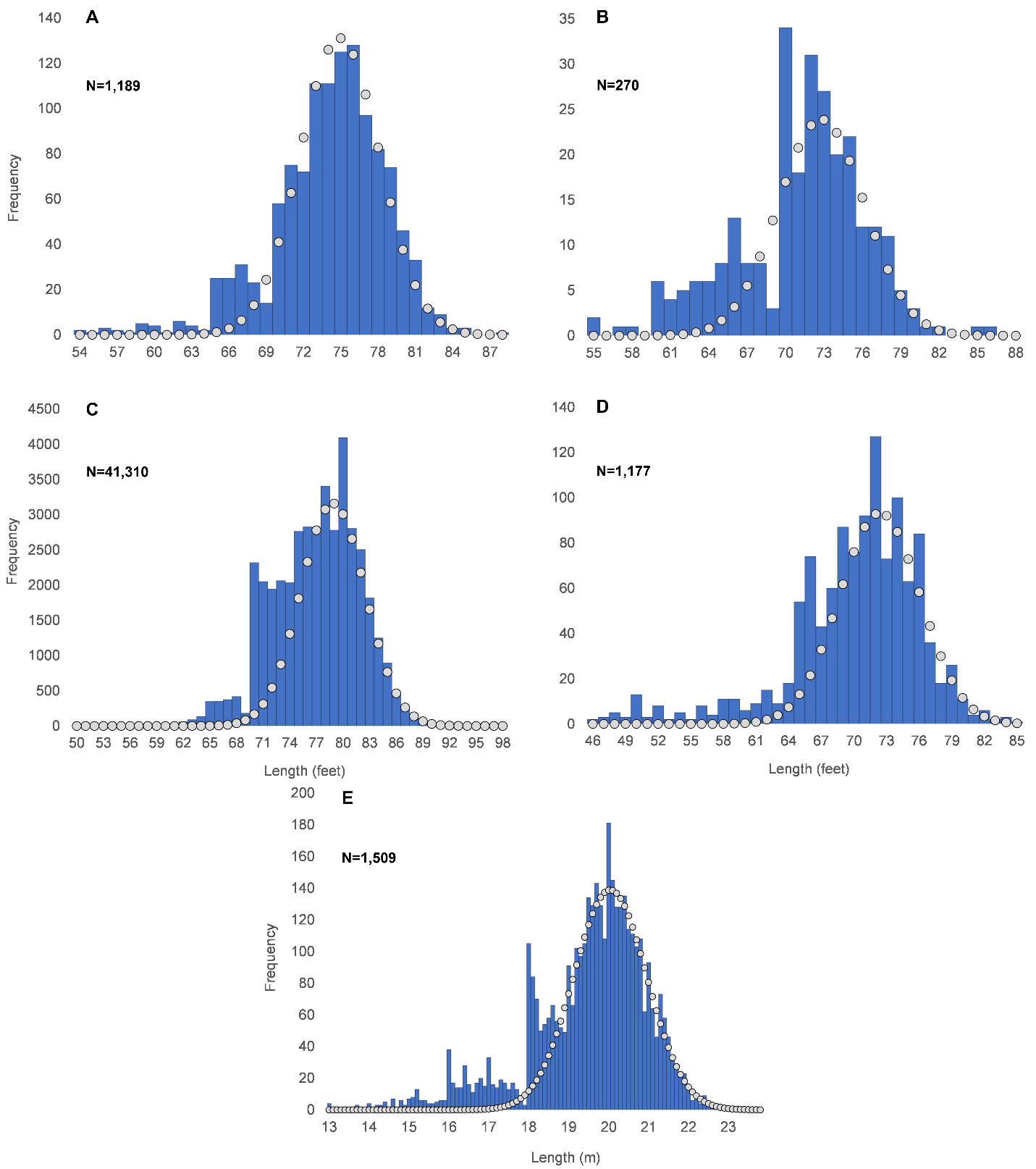

#### Figure S17. Male *B. musculus* normal probability plots for North Pacific (A), North Atlantic (B), Antarctic (C), pygmy (D), and Chilean (E).

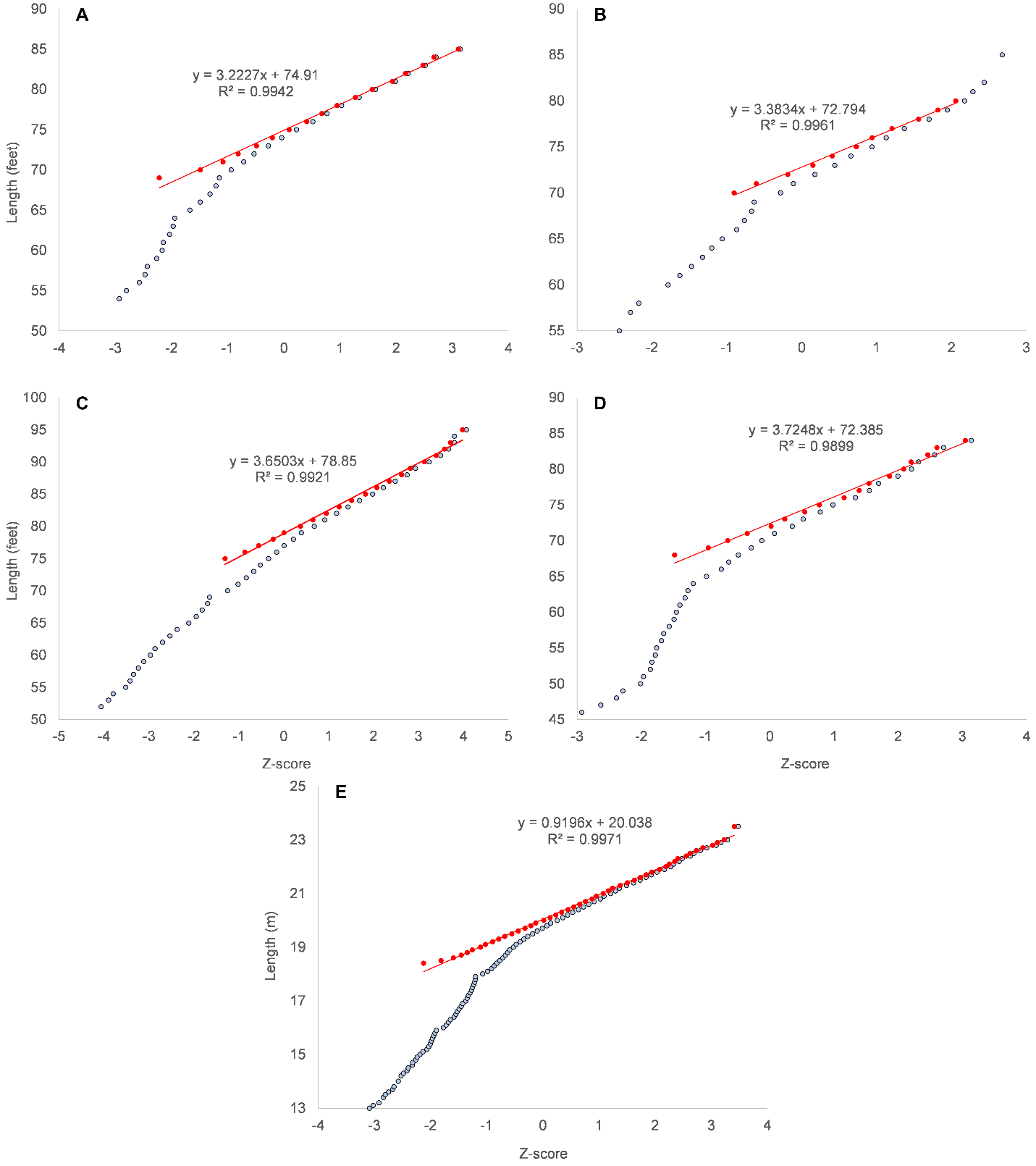

#### Figure S18. Linear weight-length regression for *M. novaeangliae* (y=2.974x-1.835, n=33, r^2^=0.961). See Supplementary File 2 for Appendix and File 3 for references.

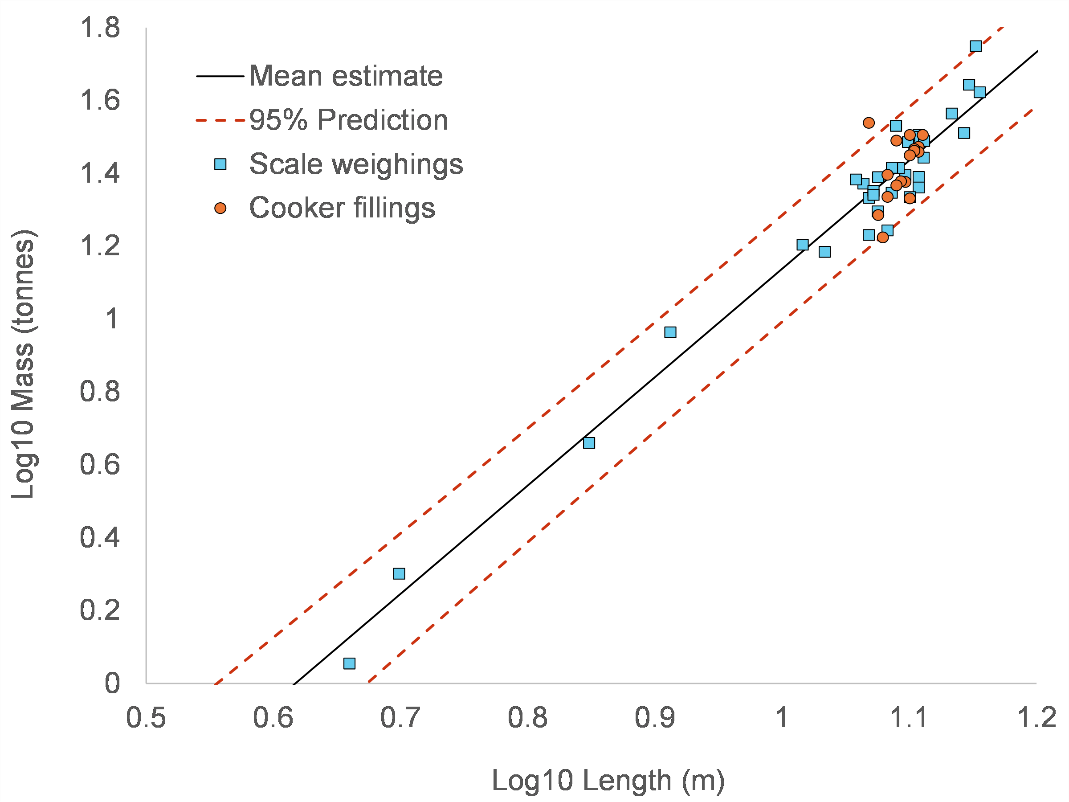

#### Figure S19. (A) Linear weight-length regression for *B. borealis* (y =3.175x– 2.342, n= 60, r^2^= 0.889). See Supplementary File 2 for Appendix and File 3 for references.

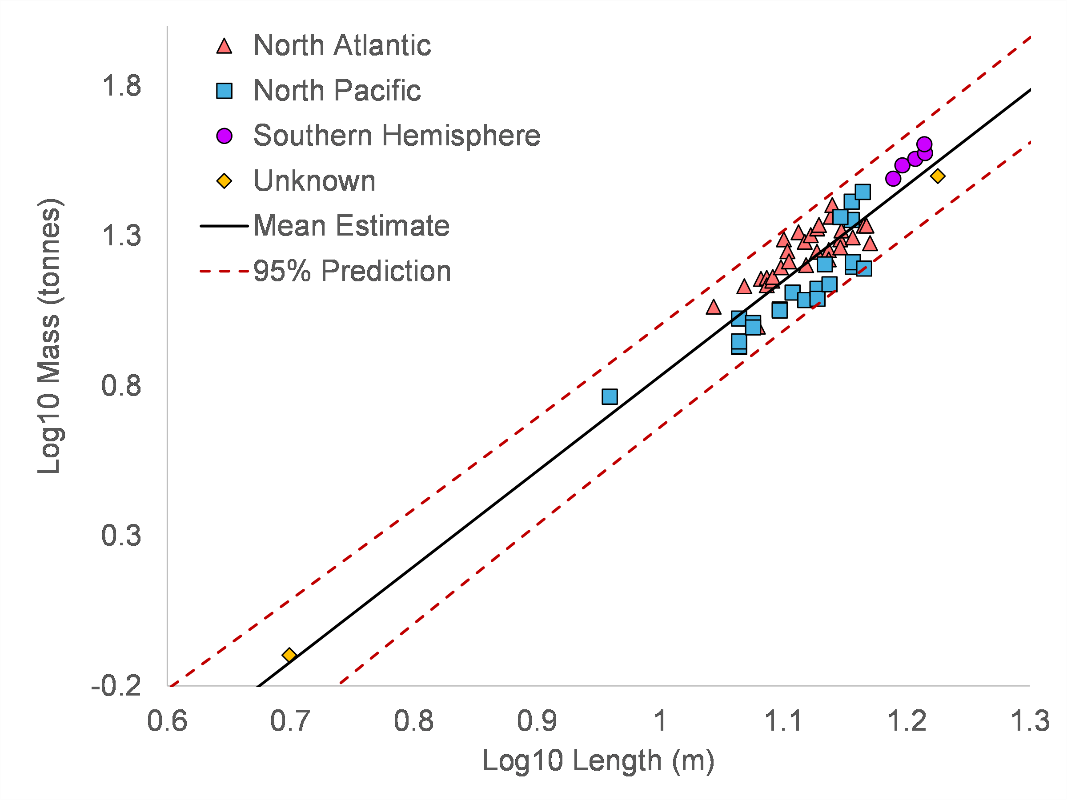

#### Figure S20. (A) Linear bone weight-body length regression for southern fin whales (y = 2.549x – 2.443, n= 40, r^2^= 0.856). (B) Normal probability plot of total weight/bone weight ratios for southern fin whales. See File 2 for appendix and File 3 for references.

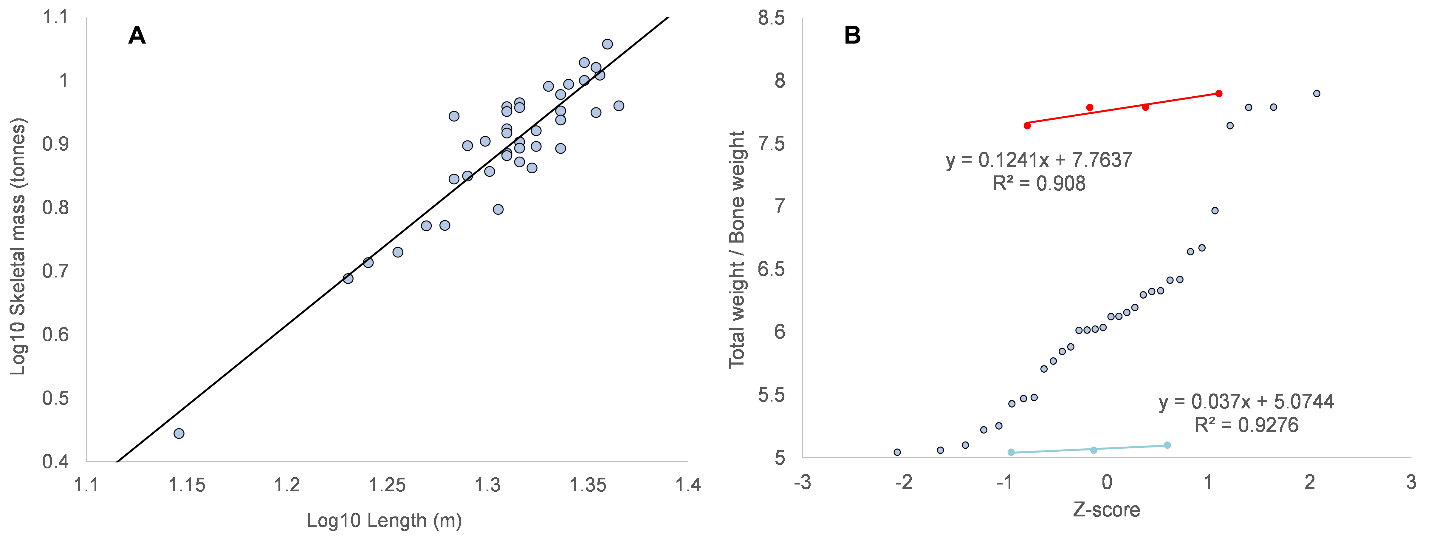

#### Figure S21. Linear bone weight-body length regression for *B. musculus* (y = 3.283x – 3.373, n= 43, r^2^= 0.910). See File 2 for appendix and File 3 for references.

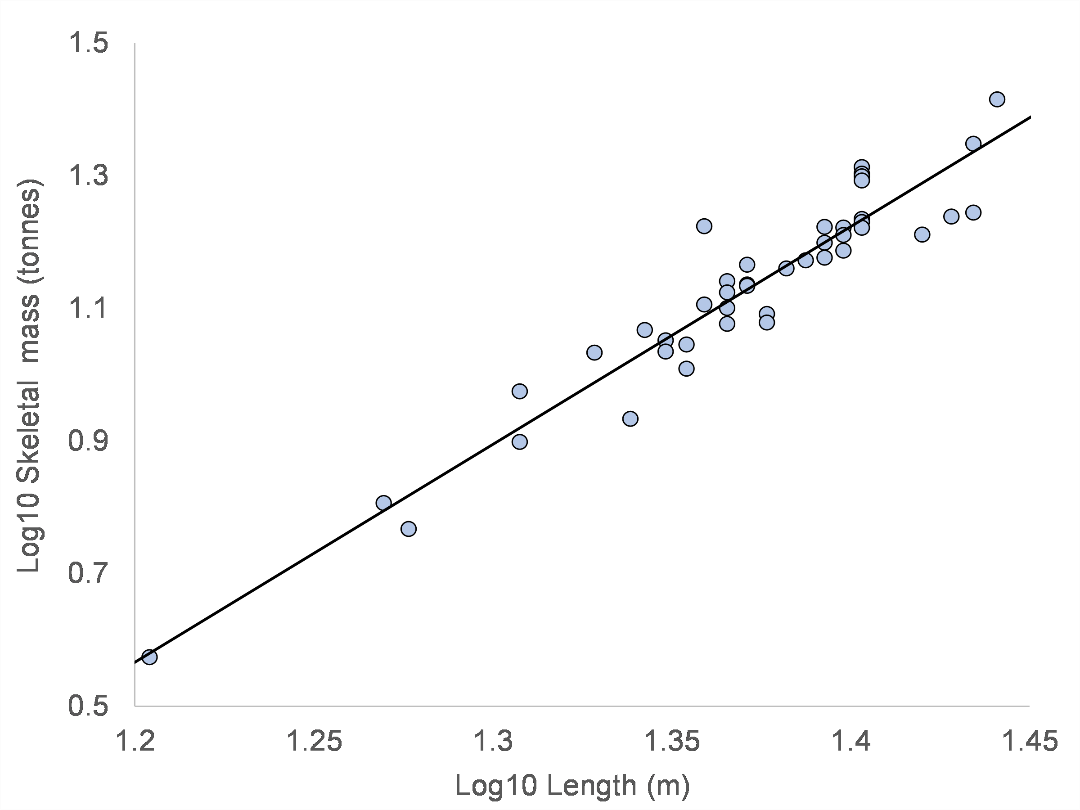

#### Table S3. Length-weight regression parameters for Balaenopteridae.

| Formula | Species | a^a^ | b/b1 | b2 | N | Reference |
| --- | --- | --- | --- | --- | --- | --- |
| Eq. 2 | *M. novaeangliae* | 0.0158 | 2.95 |  | 17 | Lockyer, 1976 |
| Eq. 2 | *M. novaeangliae* | 0.0248 | 2.794 |  | 17 | Mikhalev, 2019 |
| Eq. 2 | *M. novaeangliae* | 0.0169^b^ | 2.962 |  | 973 | Russell et al., 2023; Aoki et al., 2021 |
| Eq. 2 | *M. novaengliae* | 0.00868^b^ | 3.183 |  | 256 | Napoli et al., 2024; Aoki et al., 2021 |
| Eq. 2 | *M. novaengliae* | 0.0161 | 2.920 |  | 533 | Christiansen et al., 2025 |
| Eq. 2 | *M. novaengliae* | 0.0195 | 2.860 |  | 2,651 | van Aswegen, 2025 |
| Eq.2 | *M. novaeangliae* | 0.0192^b^ | 2.95 |  | 282 | Bernier-Graveline. 2025 |
| Eq. 2 | *M. novaeangliae* | 0.0147 | 2.970 |  | 50 | Current study |
| Eq. 2 | *B. edeni* (North Pacific) | 0.0122 | 2.74 |  | 27 | Lockyer, 1976 |
| Eq. 2 | *B. edeni* (Southern) | 0.0126 | 2.76 |  | 69 | Ohsumi, 1980 |
| Eq. 2 | *B. b. borealis* (North Pacific) | 0.0242 | 2.43 |  | 16 | Lockyer, 1976 |
| Eq. 2 | *B. b. borealis* (North Atlantic) | 0.1859 | 1.776 |  | 27 | Víkingsson et al., 1988 |
| Eq. 3 | *B. b. borealis* (North Atlantic) | 0.11279 | 1.097 | 1.324 | 27 | Víkingsson et al., 1988 |
| Eq. 2 | *B. borealis* | 0.0107 | 2.726 |  | 38 | Mikhalev, 2019 |
| Eq. 2 | *B. borealis* | 0.00455 | 3.175 |  | 60 | Current study |
| Eq. 2 | *B. p. quoyi* | 0.0238 | 2.53 |  | 43 | Lockyer, 1976 |
| Eq. 2 | *B. p. quoyi* | 0.00989 | 2.84 |  | 34 | Mikhalev, 2019 |
| Eq. 2 | *B. p. physalus* | 0.0095 | 2.865 |  | 29 | Víkingsson et al., 1988 |
| Eq. 3 | *B. p. physalus* | 0.0364 | 1.541 | 1.324 | 29 | Víkingsson et al., 1988 |
| Eq. 2 | *B. p. quoyi* (Bone mass) | 0.00360 | 2.549 |  | 40 | Current Study |
| Eq. 2 | *B. p. quoyi* (Lean) | 0.01827 | 2.549 |  | — | Current Study |
| Eq. 2 | *B. p. quoyi* (Fatten) | 0.02796 | 2.549 |  | — | Current Study |
| Eq. 2 | *B. musculus* | 0.0046 | 3.09 |  | 43 | Lockyer, 1976 |
| Eq. 2 | *B. musculus* | 0.00681 | 2.933 |  | 57 | Mikhalev, 2019 |
| Eq. 2 | *B.m. brevicauda^e^* | 0.01280 | 2.7891 |  | 43 | Russell et al., 2024 |
| Eq. 2 | *B. musculus* (Bone mass) | 0.00042 | 3.283 |  | 43 | Current study |
| Eq. 2 | *B. m. intermedia* (Lean) | 0.00197^c^ | 3.283 |  | — | Current study |
| Eq. 2 | *B. m. intermedia* (Fatten) | 0.00294^c^ | 3.283 |  | — | Current study |
| Eq. 2 | *B. m. brevicauda* (Lean) | 0.00333^d^ | 3.127 |  | — | Current study |
| Eq. 2 | *B. m. brevicauda* (Fatten) | 0.00512^d^ | 3.118 |  | — | Current study |

a. Value adjusted to meters and tonnes.
b. The proportionality constant for volume was multiplied by average tissue density of 1,037 kg / m^3^ from Aoki et al., 2021.
c. Uses tissue ratios from Lockyer, 1981
d. Uses adjust visceral component for weight.
e. Volume formula
