## Supplementary File 4 for "A summary of intraspecific size variation for large mysticetes": Supplementary File 4.R

```
########################Balaenid weight Analysis#################

#Remove outliers from balaenidmass#

balaenidmass2<-subset(balaenidmass, balaenidmass$No.!= 'EgNo2030' & balaenidmass$No.!= 'EgNo 2366' )


model1 <- lm(Log10W ~ Log10L, data = balaenidmass2)
model2 <- lm(Log10W ~ Log10L + Genus, data = balaenidmass2)
model3<- lm(Log10W ~ Log10L + Species, data = balaenidmass2)

library(AICcmodavg)

models <- list(model1, model2,model3)

mod.names <- c('Pooled', 'Separate genera','Separate Species')

aictab(cand.set = models, modnames = mod.names)

#####################Sei whale weight analysis#####################

model1 <- lm(Log10W ~ Log10L, data = seimass)
model2 <- lm(Log10W ~ Log10L + Region, data = seimass)


library(AICcmodavg)

models <- list(model1, model2)

mod.names <- c('Pooled', 'Seperate regions')

aictab(cand.set = models, modnames = mod.names)

##############################Fin whale weight analysis

###Import fin whale weight spreadsheet, after removing adjusted averages 
### highlighted in yellow and renaming to ''finmass.csv)''


model1 <- lm(log10W ~ log10L, data = finmass)
model2 <- lm(log10W ~ log10L + Location, data = finmass)

library(AICcmodavg)

models <- list(model1, model2)

mod.names <- c('Pooled regions', 'Separate regions')

aictab(cand.set = models, modnames = mod.names)

##############################Humpback whale weight analysis

###Import humpback weight spreadsheet, after renaming to ''humpbackmass'' 

model1 <- lm(log10W ~ log10L, data = humpbackmass)
model2 <- lm(log10W ~ log10L + Note, data = humpbackmass)

library(AICcmodavg)

models <- list(model1, model2)

mod.names <- c('Pooled methods', 'Separate methods')

aictab(cand.set = models, modnames = mod.names)

#############Length parameter comparisons###############

#Separating datasets between toothed and baleen whales

odontgrowth<-subset(earlycetgrowth, Clade=='Odontocete')
mystigrowth<-subset(earlycetgrowth, Clade=='Mysticete')


#comparing parameters

t_test_birth <- t.test(odontgrowth$Bfrac, mystigrowth$Bfrac)

print(t_test_birth)

#Difference significant
t_test_wean <- t.test(odontgrowth$Wfrac, mystigrowth$Wfrac)

print(t_test_wean)

########Comparing adult growth###############
#separating datasets

odontmean<-subset(lmeanpool, Clade== 'Odontocete')
mystimean<-subset(lmeanpool, Clade== 'Mysticete')

#Pooled averages
summary(lmeanpool$sm.inf)
summary(lmeanpool$mean.inf)
summary(lmeanpool$sm.mean)

#comparing adult parameters

t_test_SMLINF <- t.test(odontmean$sm.inf, mystimean$sm.inf)

print(t_test_SMLINF)

t_test_SMMEAN <- t.test(odontmean$sm.mean, mystimean$sm.mean)

print(t_test_SMMEAN)

t_test_MEANINF <- t.test(odontmean$mean.inf, mystimean$mean.inf)

print(t_test_MEANINF)

###########Correlation between CV against ratios

meanlargesamp<-subset(lmeanpool, Notes!=('Sample below 200'))

CVSMLINF<-lm(sm.inf~CV,data =meanlargesamp)

summary(CVSMLINF)

CVSMMEAN<-lm(sm.mean~CV,data =meanlargesamp)

summary(CVSMMEAN)

CVMEANINF<-lm(mean.inf~CV,data =meanlargesamp)

summary(CVMEANINF)
```
